# Parameters and determinants of responses to selection in antibody libraries

**DOI:** 10.1101/712539

**Authors:** Steven Schulz, Sébastien Boyer, Matteo Smerlak, Simona Cocco, Rémi Monasson, Clément Nizak, Olivier Rivoire

## Abstract

Antibody repertoires contain binders to nearly any target antigen. The sequences of these antibodies differ mostly at few sites located on the surface of a scaffold that itself consists of much less varied amino acids. What is the impact of this scaffold on the response to selection of a repertoire? To gauge this impact, we carried out quantitative phage display experiments with three antibody libraries based on distinct scaffolds harboring the same diversity at randomized sites, which we selected for binding to four arbitrary targets. We first show that the response to selection of an antibody library is captured by a simple and measurable parameter with direct physical and information-theoretic interpretations. Second, we identify a major determinant of this parameter which is encoded in the scaffold, its degree of evolutionary maturation. Antibodies undergo an accelerated evolutionary process, called affinity maturation, to improve their affinity to a given target antigen as part of the adaptive immune response. We find that libraries of antibodies built around such maturated scaffolds have a lower response to selection to other arbitrary targets than libraries built around naïve scaffolds of germline origin. Our results are a first step towards quantifying and controlling the evolutionary potential of biomolecules.

## 1. Introduction

The idea that evolution by natural selection is not only leading to adaptations but to a propensity to adapt, or “evolvability”, has been repeatedly put forward [1, 2, 3]. As demonstrated by a number of mathematical models, evolvability can indeed emerge from evolutionary dynamics without any direct selection for it [4, 5, 6, 7]. Yet, theoretical insights have not translated into experimental assays for measuring and controlling evolvability in actual biological systems. Biomolecules as RNAs and proteins are ideal model systems for developing such assays as they are amenable to controlled experimental evolution [8]. For proteins, in particular, several biophysical and structural features have been proposed to correlate with their evolvability, most notably their thermal stability [9] and the modularity and polarity of their native fold [10]. A major limitation, however, is the absence of a measurable index of evolvability quantifying evolutionary responses to compare to biophysical or structural quantities.

Here, we present results of quantitative selection experiments with antibodies that address this issue. Antibodies are particularly well suited to devise and test new approaches to measure and control evolvability. They conveniently span a large phenotypic diversity, specific binding to virtually any molecular target, by means of a limited genotypic diversity. Most of the diversity of natural antibody repertoires is indeed achieved by a few randomized loops that are displayed onto a structurally more conserved framework [11]. Further, well-established screening techniques are available for manipulating libraries of billions of diverse antibodies [12]. More fundamentally, antibodies are subject to two evolutionary processes on two distinct time scales: their frameworks evolve on the time scale of many generations of their host, as all other genes, and both frameworks and loops also evolve on a much shorter time scale as part of the immune response in the process of affinity maturation [13]. Importantly, affinity maturation-associated mutations are somatic and the sequences of maturated antibodies are not transmitted to subsequent generations. Germline antibody frameworks, whose transmitted sequences are the starting point of affinity maturation, are thus well positioned to be particularly evolvable, as evolving to increase their affinity to antigens is part of their physiological role.

As a first step towards quantifying and controlling the evolvability of antibodies, we previously characterized the response to selection of antibody libraries built around different structural scaffolds [14]. We took for these scaffolds the frameworks of heavy chains (V_H_) of natural antibodies, and built libraries by introducing all combinations of amino acids at four consecutive sites in their complementary determining region (CDR3) loop, a part of their sequence known to determine their binding affinity and specificity [11]. Using phage display [15], we selected sequences from these libraries for their ability to bind different molecular targets and inferred the relative enrichment, or selectivity, of different antibody sequences by high-throughput sequencing [16]. Comparing experiments with libraries built on different scaffolds and selected against different targets led us to two conclusions. First, we quantified the variability of responses to selection of different sequences within a library and found this variability to differ widely across experiments involving different libraries and/or different targets. Second, we observed a hierarchy of selectivities between libraries, with multiple sequences from one particular library dominating selections involving a mixture of different libraries. These results raised two questions: (i) How to relate the hierarchies of selectivities between and within libraries? (ii) How to rationalize the differences between scaffolds that are all homologous?

Here, we answer these two questions through the presentation of new data and new analyses. First, we propose to characterize the hierarchies within and between libraries with two parameters for which we provide interpretations from the three standpoints of physics, information theory and sequence content. Second, we present new experimental results that identify the degree of maturation of an antibody scaffold as a control parameter for its selective potential. The results are, to our knowledge, the first demonstration based on quantifying the evolutionary responses to multiple selective pressures that long-term evolution has endowed germline antibody frameworks with a special ability to respond to selection.

## 2. Methods

### 2.1. Experimental design

In the absence of mutations, the outcome of an evolutionary process is determined by the properties of its initial population. In our experiments where antibodies are evolved in successive cycles of selection and amplification, the critical property of a sequence *x* present in the initial population is its selectivity *s*(*x*), the factor by which it is enriched or depleted from one cycle to the next (see Box). Selection involves binding to a target, which is varied between experiments. Experiments are designed for the selectivity *s*(*x*) to reflect the binding affinity of sequence *x* to this target (see Appendix 1.1). Inevitably, however, it can also depend on affinity to non-target substrates and to sequence-dependent differences in amplification. Importantly, while the details of these “biases” are contingent to the experimental approach, their presence is a generic feature of any process of molecular evolution, including the natural process of antibody affinity maturation of which the experiments mimic the first step, prior to the introduction of any mutation.

Each of our libraries consists of sequences with a common part, which we call a scaffold, and 4 positions that are randomized to all *N* = 20^4^ combinations, where 20 is the number of natural amino acids. The mapping *x* ↦ *s*_*L,T*_ (*x*) from 4-position sequences *x* to selectivities thus depends both on the scaffold that defines the library *L* and on the target *T* that defines the selective pressure. We are interested in properties of the scaffold that favor large values of selectivities, where “large” is considered either relative to other sequences within the same library (same scaffold) or relative to sequences from different libraries (different scaffolds).

Our previous experiments involved 24 different libraries, each built on a different scaffold consisting of a natural V_H_ fragment [14]. These fragments originate from the germline or the B cells of organisms of various species. Scaffolds from the germline encode naïve antibodies which have not been subject to any affinity maturation, while scaffolds from B cells encode maturated antibodies which have evolved from naïve antibodies to bind strongly to antigens encountered by the organisms. We previously performed experiments where the initial population consisted either of a single library or a mixture of different libraries [14]. In particular, in two experiments using very different targets (a neutral polymer and a DNA loop) we coselected all 24 libraries together. Strikingly, while only 2 of the 24 libraries were built on germline scaffolds, the final population of one experiment was dominated by antibodies built on one of the two naïve scaffolds, and the second by the other one. This suggested us that germline scaffolds may have an intrinsically higher selective potential.

To investigate this hypothesis, we analyze here the selection against 4 different targets of 3 libraries with varying degrees of maturation. The scaffolds of the 3 libraries originate from Human V_H_ fragments and have evolved to different degrees as part of the immune response of patients infected by HIV (Fig. S1). The first scaffold (Germ) is taken from the germline and has not undergone any maturation. The second scaffold (Lim) has been subject to limited affinity maturation and differs from Germ, from which it originates, by 14 % of its amino acids. The third scaffold (Bnab) is a so-called broadly neutralizing antibody [17], which has evolved over many years to recognize a conserved part of the HIV virus [18]; it also originates from Germ, to which it differs by 34 % of its amino acids, and has evolved independently of Lim, to which it differs by 38 %. The 3 libraries, which are built around these scaffolds by introducing all combinations of amino acids at 4 positions of their CDR3 were part of the 24 libraries used in our previous experiments [14]. Here, to systematically compare the selective potential of these libraries, we present experiments where they are selected against four different targets, two DNA targets (DNA hairpins with a common stem but different loops, denoted DNA1 and DNA2, Fig. S2) and two homologous protein targets (the fluorescent proteins eGFP and mCherry, denoted prot1 and prot2), each unrelated to the HIV virus against which the Lim and Bnab scaffolds had been maturated.

### 2.2. Parametrization

To quantitatively compare the outcome of different experiments with different libraries and targets, we introduce here two parameters, *σ* and *µ*, which respectively quantify intra and inter-library differences in selectivities. These parameters derive from a statistical approach that considers only the distribution *P* (*s*) of values that selectivities take across the different sequences of a library [19, 20, 21]. They correspond to the assumption that this distribution is log-normal,

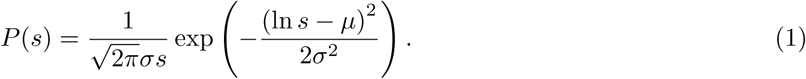

The parameter *σ* captures intra-library differences in response to selection while the parameter *µ* provides the additional information required to describe inter-library differences.

The assumption that distributions of selectivities are log-normal has several justifications. First, it empirically provides a good fit of the data, not only in our experiments as we show below, but in a number of previous studies of antibody-antigen interactions [22] and protein-DNA interactions [23], including studies that had access to the complete distribution *P* (*s*) [23]. Second, log-normal distributions are stable upon iteration of the evolutionary process: if two successive selections are performed so that *s* = *s*_1_*s*_2_ with *s*_1_ and *s*_2_ independently described by log-normal distributions with parameters (*σ*_1_, *µ*_1_) and (*σ*_2_, *µ*_2_), *s* also follows a log-normal distribution with parameters 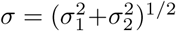 and 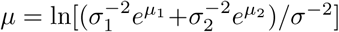; more generally, log-normal distributions are attractors of evolutionary dynamics [24]. Third, log-normal distributions are physically justified from the simplest model of interaction, an additive model where the interaction energy between sequence *x* = (*x*_1_, *…*, *x*_*𝓁*_) of length *𝓁* and its target takes the form 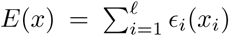 with contributions *ϵ*_*i*_(*x*_*i*_) from each position *i* and amino acid *x*_*i*_, and thus its selectivity 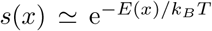, where *T* is the temperature and *k*_*B*_ the Boltzmann constant (Appendix 1.1). At thermal equilibrium and for sufficiently large *𝓁*, a log-normal distribution of the affinities is then expected with *µ ∼* −*𝓁*⟨*ϵ*⟩*/k*_*B*_*T* and*σ ∼ 𝓁*^1*/*2^(⟨*ϵ*^2^⟩ − ⟨*ϵ*⟩_2_xs)^1*/*2^*/k*_*B*_*T*, where ⟨*ϵ*⟩ and ⟨*ϵ*^2^⟩ − ⟨*ϵ*⟩_2_ are respectively the mean and variance of the values of binding energies per position *ϵ*_*i*_(*x*_*i*_).

### 2.3. Inference of parameters

Selectivities are measured as relative enrichments of sequences in two successive rounds of selection. We obtain the parameters *σ* and *µ* by fitting the values with truncated log-normal distributions (Fig. 1A and Appendix 3.3). This inference is complicated by two factors: only the upper tail of the distribution of selectivities is sampled in the experiments and enrichments provide selectivities only up to a multiplicative factor (see Box). While the parameter *σ* is independent of this multiplicative factor, comparing the parameters *µ* between libraries requires performing selections where different libraries are mixed in the initial population. To refine and validate our inference, we also performed selection experiments where we mixed a very small number of random and top selectivity sequences (Fig. 1B): as the random sequences typically reflect the mode of the distributions, (the most likely selectivity value), these experiments provide an independent estimation of *µ* that we can profitably use (see details in Appendix 3.3).

**Figure 1:**
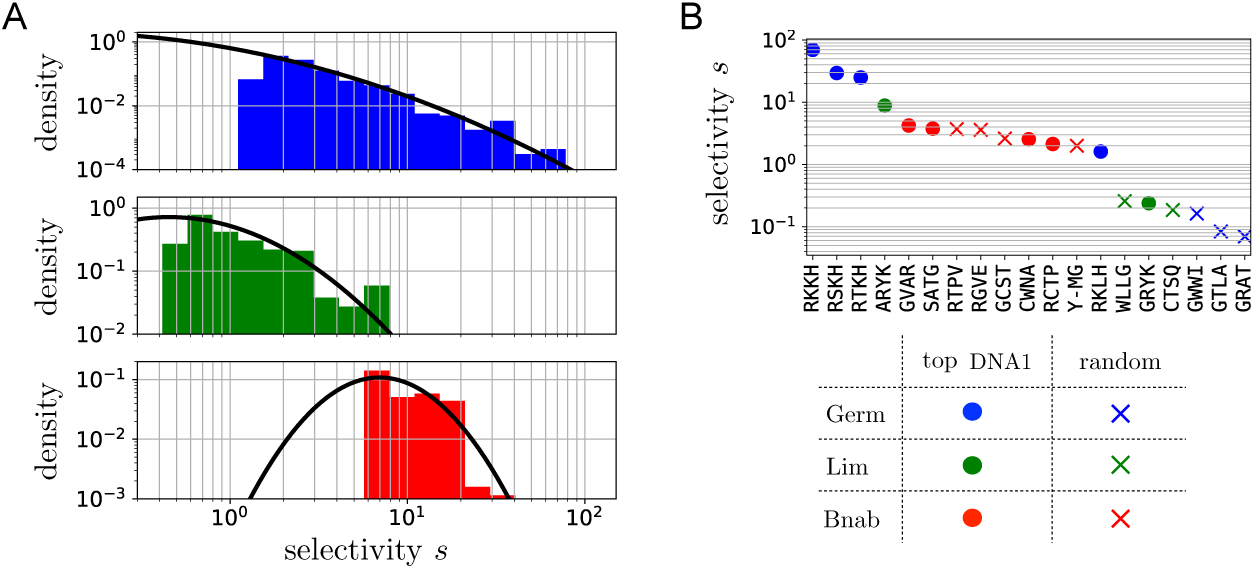
Fitting empirical distributions of selectivities with log-normal distributions. **A.** The selection of a library *L* against a target *T* provides the selectivities of the sequences in *L* that have top selectivity against *T*. Here, the histograms show the selectivities obtained from experiments where the Germ (in blue), Lim (in green) and Bnab (in red) libraries were selected against the DNA1 target. The black line is the best fit to a log-normal distribution. **B.** To locate precisely the mode of the distributions, we performed experiments where the initial population consists in a mixture of very few top (dots) and random (crosses) sequences. Top sequences are identified from A based on the largest selectivities against the target. Random sequences, on the other hand, are picked at random in the libraries and are expected to have typical selectivities located at the maxima of the black curves in A. Taken together, the results indicate that when selected against the DNA1 target, the Germ library has the highest *σ* and the Bnab library the highest *µ*. Similar results are obtained for selections against other targets (Fig. S8 and Table 1).

The values of *σ* and *µ* that we infer for the 3 libraries Germ, Lim and Bnab when selected against each of the 4 targets DNA1, DNA2, prot1 and prot2 are presented in Fig. 2A. We validated the quality of the fits by probability-probability and quantile-quantile plots (Figs. S16-S18). We also assessed the robustness of the inference by comparing replicate experiments (Figs. S16-S22), and comparing experiments where a library is selected either alone or in mixture with the other two (Fig. S19). Finally, we verified that the results are unchanged whether selectivities are measured from enrichments between the 2nd and 3rd cycles, or the 3rd and 4th ones (Figs. S20-S21).

**Table 1:**
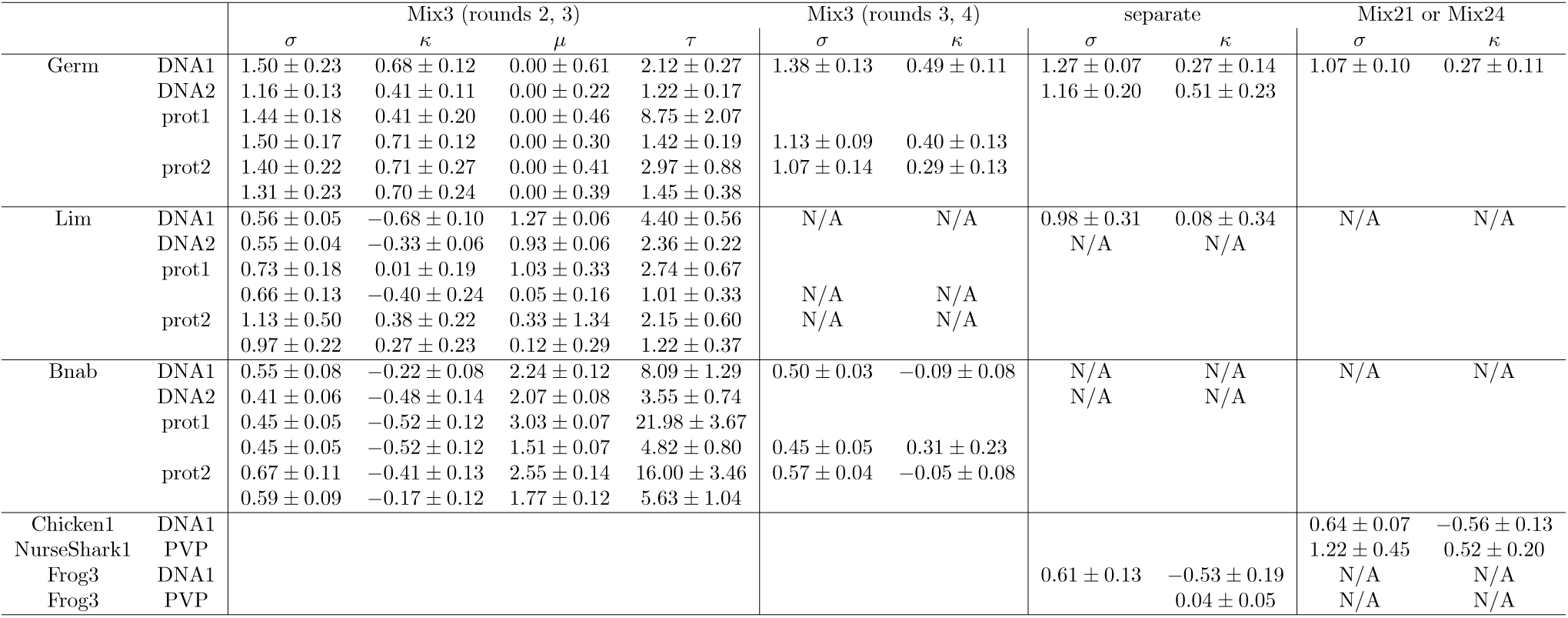
Parameters obtained from fits of the distribution of selectivities to generalized Pareto distributions (*κ, τ*) and log-normal distributions (*σ, µ*) for experiments presented here and in our previous work [14]. N/A indicates that the data was insufficient to make a meaningful fit. For selectivities against the protein targets between rounds *c* = 2 and *c* + 1 = 3, values are given for two independent replica of the experiment. The given uncertainties correspond to a single standard deviation around the maximum likelihood estimate as given by the Cramer-Rao bound. In the case of Frog3 against DNA1, and only in this case, the value of *κ* differs from the one reported in our previous work [14] for reasons explained in Appendix 3.2 and Figure S15.

**Figure 2:**
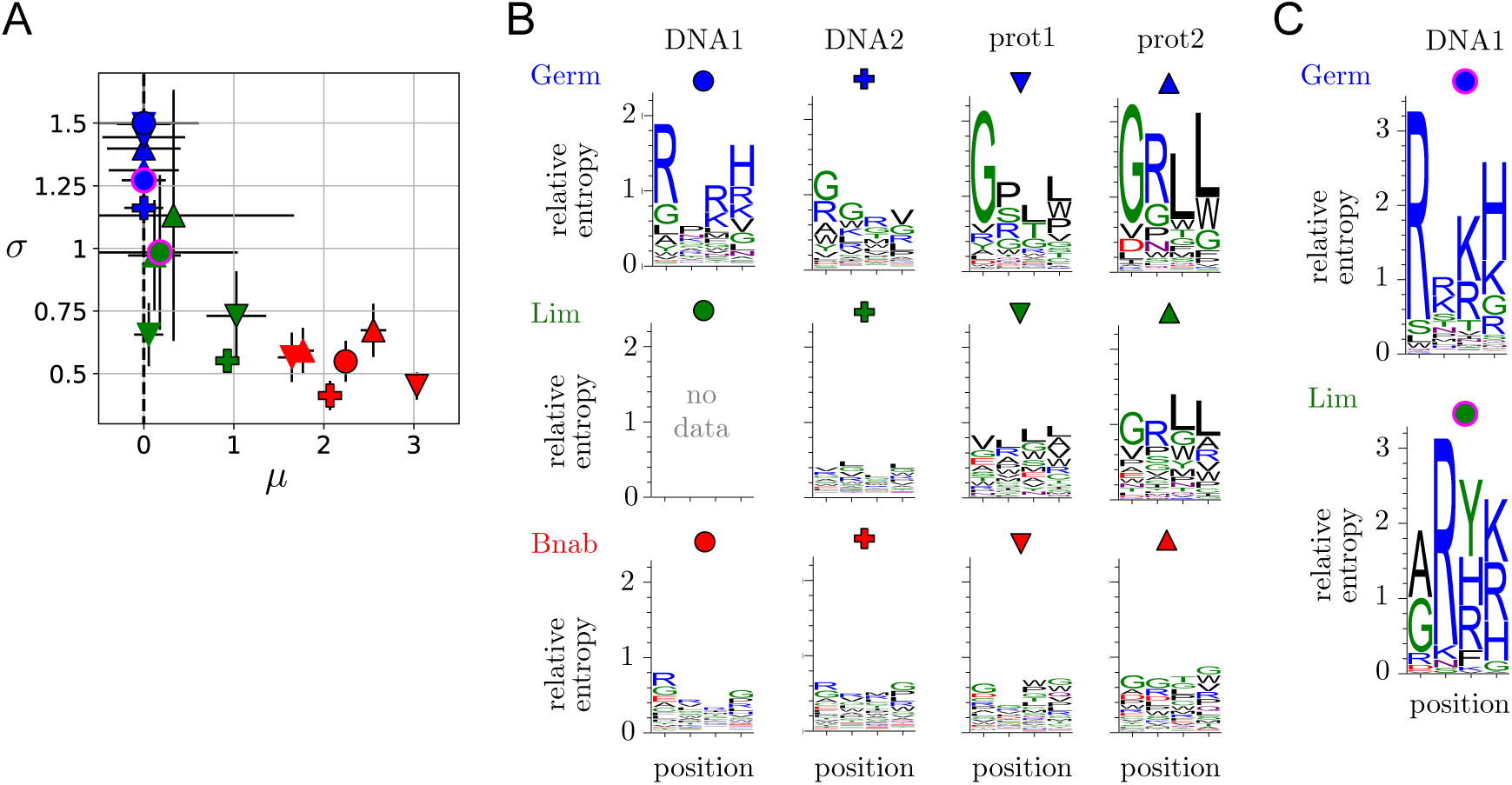
Comparing selections of libraries built on scaffolds with different degrees of maturation – **A.** Parameters (*µ, σ*) of the distributions of selectivities for our 3 libraries selected against 4 targets. The color of the symbols indicates the library (Germ, Lim or Bnab) and its shape the target (DNA1, DNA2, prot1 or prot2) with the conventions defined in B. Symbols with a black or no contour indicate results from replicate experiments where the 3 libraries are mixed in the initial population, and symbols with a magenta contour where a library is screened in isolation. *µ*_Germ,*T*_ is conventionally set to *µ*_Germ,*T*_ = 0 for all targets *T* (Appendix 3.4). *µ* is generally more challenging to infer than *σ* and it shows here more variations across replicate experiments. **B.** Sequence logos for 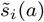, which represent the contribution of the different amino acids to the selectivities (see Box), for the selections of the three libraries, Germ, Lim and Bnab against the two DNA targets (DNA1 and DNA2) and the two protein targets (prot1 and prot2). These results correspond to experiments where the 3 libraries are mixed in the initial population. The Lim library is outcompeted by the other two libraries when selected against the DNA1 target, which does not leave enough sequences to make a meaningful inference (see also Fig. S10 for more details on the sequence logos for the Bnab library). **C.** Sequence logos for 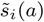 for the Germ and Lim libraries selected in isolation against the DNA1 target. For the Lim library, this palliates the absence of data in B. For the Germ library, it shows that essentially the same motifs are found whether the library is selected in a mixture as in B or on its own; the area under the logos is, however, differenϵ it would be *σ*^2^*/*2 with infinite sampling, but major deviations are caused by limited sampling (Fig. S9).

## 3. Results

### 3.1. Intra-library hierarchy

The hierarchy of selectivities within a library is quantified by the parameter *σ*: a small *σ* indicates that all sequences in the library are equally selected while a large *σ* indicates that the response to selection varies widely between sequences in the library. When comparing the *σ*_*L,T*_ inferred from the selections of the 3 libraries *L* against each of the 4 targets *T*, a remarkable pattern emerges: the more a scaffold is maturated, the smaller is *σ, σ*_Germ,*T*_ *> σ*_Lim,*T*_ ≥ *σ*_Bnab,*T*_ for all targets *T*, and even min_*T*_ (*σ*_Germ,*T*_) *>* max_*T*_ (*σ*_Lim,*T*_, *σ*_Bnab,*T*_) (Fig. 2A). Statistically, if considering the inequalities to be strict, the experiments to be independent and any result to be *a priori* equally likely, the probability of this finding is only *p* = (3!)^−4^ ≃ 7.10^−4^.

Examining sequence logos shows that although selections of the Germ library are characterized by a similarly high value of *σ* for the 4 targets, the sequences that are selected against each target are different (Fig. 2B-C). The amino acids found to be enriched are consistent with the nature of the targets: selections against the DNA targets are dominated by positively charged amino acids (letters in blue) and selections against the two protein targets, which are close homologs, are dominated by similar amino acid motifs.

In contrast, sequences logos for the Bnab library show motifs that are less dependent on the target (Fig. 2B and Fig. S10). This observation is rationalized by an experiment where only the amplification step is performed, in the absence of any selection for binding. Sequence-specific amplification biases are then revealed, with sequence motifs that are similar to those observed when selection for binding is present (Fig. S10). With protein targets at least, the motifs are nevertheless sufficiently different to infer that selection for binding to the target contributes significantly to the selectivities (see also Fig. S6). Target-specific selection for binding, which is dominating the top selectivities in the Germ library (Fig. S11), is thus of the same order of magnitude as amplification biases for the top selectivities in the Bnab library.

Remarkably, the Lim library behaves either like the Germ library or the BnAb library, depending on the target. In particular, a motif of positively charged amino acids emerges when selecting it against one of the two DNA targets (DNA1), but no clear motif emerges when selecting it against the other one (DNA2). Besides, when a clear motif emerges, it can be identical to the motif emerging from the Germ library as in case of a selection against the prot2 target, or different, as in the case of a selection against the DNA1 target (but with a similar selection of positively charged amino acids).

### 3.2. Inter-library hierarchy

The hierarchy of selectivities between libraries is quantified by the parameter *µ*. This parameter also shows a pattern that is independent of the targeϵ *µ*_Germ,*T*_ ≃ *µ*_Lim,*T*_ *< µ*_Bnab,*T*_ and even max_*T*_ (*µ*_Germ,*T*_, *µ*_Lim,*T*_) *<* min_*T*_ (*µ*_Bnab,*T*_) (Fig. 2B). Inferring *µ* is more challenging than inferring *σ* and the differences observed between the Germ and Lim libraries are most likely not significant, as apparent from the observed variations between replicate experiments. The *µ* of the Bnab library is, on the other hand, systematically larger. The difference is explained by an experiment where selection is performed in the absence of DNA or protein targets but in the presence of streptavidin-coated magnetic beads to which these targets are usually attached.

This experiment reproduces the differences in *µ*_*L,T*_, which indicates a small but significant affinity of the Bnab scaffold for the magnetic beads, independent of the sequence *x* (Fig. S12). While the differences in *σ* appear to be independent of the target, the differences in *µ* are thus related to a common feature of the targets. Given these different origins, the correlation between *σ* and *µ* that we observe may be fortuitous.

### 3.3. Implications for evolutionary dynamics

The different patterns of intra and inter-library hierarchies lead to a non-trivial evolutionary dynamics when selecting from an initial population that is composed of different libraries. In particular, a non-monotonic enrichment is expected when mixing two libraries characterized by (*µ*_1_, *σ*_1_) and (*µ*_2_, *σ*_2_) with *µ*_1_ *> µ*_2_ but *σ*_1_ *< σ*_2_: the library with largest *µ* dominates the first cycles while the one with largest *σ* dominates the later ones. This is indeed observed in experiments where different libraries are mixed in the initial population (Fig. 3). The dynamics of the relative frequencies of different libraries is globally predicted by a calculation of library frequencies in the mix based on the parameters (*µ, σ*) inferred for each library independently (Appendix 1.4), even though deviations are expected from the non-uniform sampling of sequences within each library, which is not encoded in *σ* or *µ*. Parametrizing the response to selection of a library by the two parameters (*µ,σ*) is thus not only useful to characterize its intrinsic response but also to rationalize the evolutionary dynamics of mixtures of libraries.

**Figure 3:**
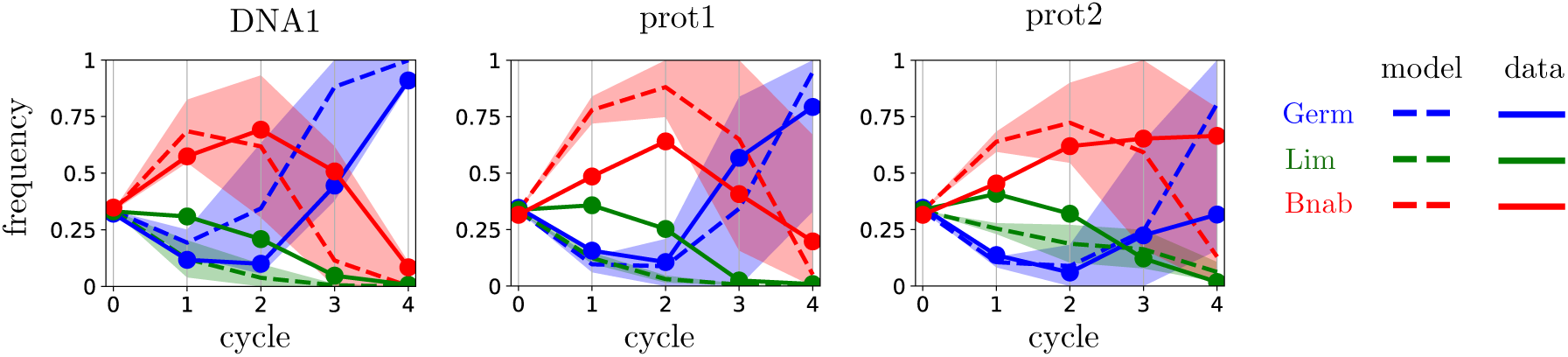
Dynamics of library frequencies – A mixture of the three libraries, Germ (blue), Lim (green) and Bnab (red) was subject to four successive cycles of selection and amplification against different targets. The full lines report the evolution of the relative frequencies of the three scaffolds. The dotted lines represent the estimated dynamics using the characterization of each library by a log-normal distribution with the parameters *σ, µ* estimated from the selection of the libraries against the same target (Appendix 1.4). The shaded area correspond to one standard deviation in the estimation of the parameters *σ, µ*. The model assumes that sequences are uniformly represented in each initial library, which is not the case in experiments and explains why the agreement with the data is only qualitative.

## 4. Discussion

How to interpret the result that intra-library diversity, as characterized by *σ*, decreases with the level of maturation of the scaffold? Here, we show that the parameter *σ* provides a characterization of intra-library diversity that is equivalent to three other approaches based on extreme value theory, information theory and sequence logos.

### 4.1. Extreme value statistics

In our previous work [14], we fitted the tail of the distribution of selectivities with generalized Pareto distributions, a family of distributions with two parameters, a shape parameter *κ* and a scaling parameter *τ*. This was motivated by extreme value theory, which establishes that these parameters are sufficient to describe the tail of any distribution (Appendix 1.2). For different libraries *L* and different targets *T*, we found that generalized Pareto distributions provide a good fit of the upper tail of *P*_*L,T*_ (*s*), with, depending on the scaffold *L* and target *T* either *κ >* 0 (heavy tail), *κ <* 0 (bounded tail) or *κ* = 0 (exponential tail). The origin of these different values of *κ* was, however, unclear.

Comparing probability-probability plots to assess the quality of the fits, our data appears equally well fitted by generalized Pareto distributions and log-normal distributions (Figs. S16-S22). This finding is at first sight puzzling as some of the fits with generalized Pareto distributions involve a non-zero shape parameter *κ* ≠ 0 but extreme value theory states that the tail of log-normal distributions is asymptotically described by a shape parameter *κ* = 0 for all values of *σ, µ* [25]. Extreme value theory is, however, only valid in the double asymptotic limit *N* → ∞ and *s*\* → ∞, where *N* is the total number of samples and *s*\* the threshold above which these samples are considered. With finite data, determining whether this asymptotic regime is reached is notoriously difficult when the underlying distribution is log-normal [26]. More precisely, *N* points randomly sampled from a log-normal distribution with parameter *σ* are known to display an apparent *κ*_*N*_ = *σ/*(2 ln *N*)^1*/*2^ which tends to zero only very slowly with increasing values of *N* [26]. In fact, this relationship itself requires *N* (or *σ*) to be sufficiently large and finite size effects can even produce an apparent *κ*_*N*_ *<* 0 (Fig. S14).

While casting doubt on the practical applicability of extreme value theory, these statistical effects do not call into question the main conclusion of our previous work [14]: different combinations of scaffolds *L* and targets *T* exhibit different within-library hierarchies, which are quantified by the different values of their (apparent) shape parameter *κ*. Fits with a log-normal distribution provide another parameter *σ* that report essentially the same differences (Fig. 4). More importantly, we verify on our previous data, which partly involves different scaffolds and different targets, that libraries built on germline scaffolds have a higher *σ* than libraries built around maturated scaffolds (Fig. 4 and Table 1).

**Figure 4:**
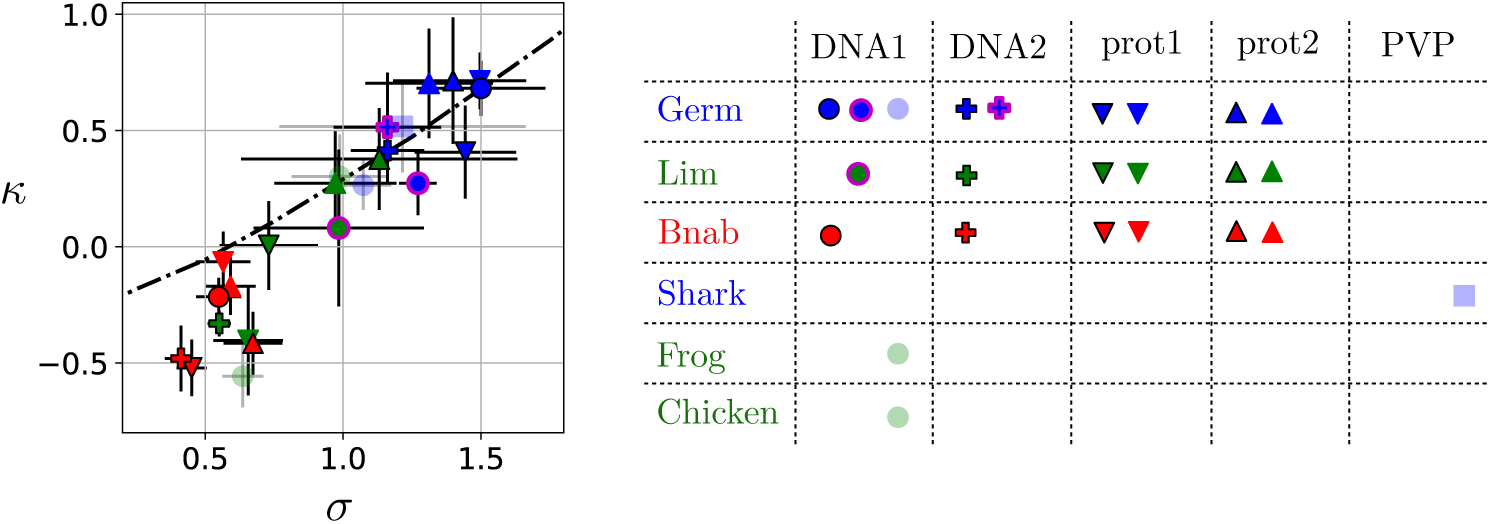
Shape parameter *κ* from fits of the selectivities to generalized Pareto distributions versus *σ* from fits to log-normal distributions – Results from different libraries selected against different targets are represented here with the same convention as in Figure 2: blue, green and red plain colors for the Germ, Lim and Bnab libraries, circle, cross, downward and upward triangles for the DNA1, DNA2, prot1 and prot2 targets. In addition, results from our previous work [14] are indicated in transparent blue if they involve a library built onto a germline scaffold and in transparent green if they involve a library built onto a maturated scaffold. The hierarchy indicated by *κ* is essentially the same as the hierarchy indicated by *σ*, consistent with the expected relationship between *κ* and *σ* (black dotted line, Fig. S14). By the two approaches, libraries built onto germline scaffolds are found to have a more diverse response to selection than libraries built onto maturated scaffolds irrespectively of the target (all values of *σ* and *κ* are given in Table 1).

### 4.2. Informational interpretation

The parameter *σ* can also be given an information-theoretic interpretation. From a statistical standpoint, the specificity of selection of a population is naturally quantified by the relative entropy *D*(*f* ^1^ ‖*f* ^0^) = Σ*x f* ^1^(*x*) ln *f* ^1^(*x*)*/f* ^0^(*x*) which compares the distribution *f* ^1^(*x*) of sequences *x* after one cycle of selection-amplification to their initial distribution *f* ^0^(*x*). When taking this initial distribution *f* ^0^(*x*) to be a uniform distribution over the *N* possible sequences, *f* ^0^(*x*) = *N* ^−1^, *f* ^1^(*x*) is nothing but the selectivity 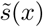 obtained by choosing *λ* in Eq. (4) such that 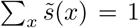. The inverse of 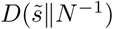 answers the following statistical question (Appendix 1.3): how large should the initial population be to infer from the outcome of an experiment that selection is at work? Assuming a large initial library with selectivity distribution *P* (*s*), 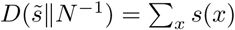 ln [*S(x)N*] can be rewritten as

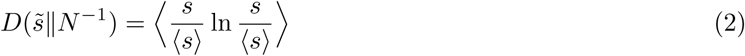

where the average is taken with *P* (*s*), i.e.,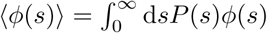. When *P* (*s*) is log-normal,

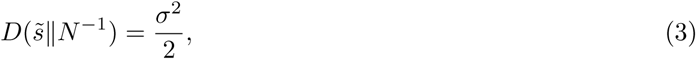

showing that *σ* can be interpreted as quantifying the specificity of selection at the population level. This statistical viewpoint can be extended to characterize the specificity of arbitrary sets of binders and ligands (Appendix 1.3), generalizing a proposal to define specificity as an amount of information encoded in interactions [27].

### 4.3. Sequence motifs

Assuming that the different sites *i* along the sequence contribute independently to the selectivity, 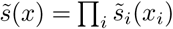, the specificity 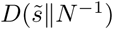 is nothing but 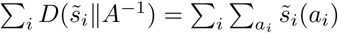 ln 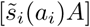, the total area under the sequence logos of 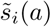, where *A* = 20 is the total number of amino acids. By displaying both amino acid specificities and an overall measure of specificity of selection 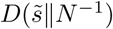, sequence logos thus provide a convenient summary of selection within a library.

This comes, however, with an important caveat when selectivities are available only for a small subset of *N* ′ ≪ *N* sequences, as it is the case in experiments. If ignoring unobserved sequences when computing 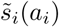, the empirically determined quantity 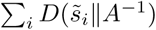 overestimates the true value of 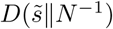, all the more as *N* ′ is smaller (Fig. S9). Because of this effect, the areas under the curve of the sequence logos based on 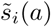 are not comparable to *σ*^2^*/*2 as Eq. (3) would suggest. They are also not comparable across different experiments when the sampling sizes *N* ′ differ (Fig. 2B and C). Finally, even with *N* ′ = *N*, deviations between 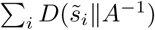 and 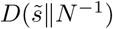 may arise if the contributions of the different positions are not additive.

## 5. Conclusion

In summary, we find that libraries built around germline antibody scaffolds have a response to selection that is quantitatively different from libraries built around maturated scaffolds: for arbitrary targets, they contain sequences with a wider range of affinities, including specific sequences with the strongest affinities. This constitutes the first quantitative evidence that germline antibodies are endowed with a special evolutionary ability to generate selectable diversity. Our work was centered onto 3 libraries, one based on a germline scaffold and two based on scaffolds derived from this germline scaffold with different degrees of maturation, which we selected against 4 different targets. Assessing the generality of our conclusion will require further experiments with additional scaffolds and targets. The statistical framework that we introduced here provides the required tools to perform this analysis systematically and quantitatively. Beyond the 3 libraries studied here, our conclusions are also supported by our previous results [14], which involved a library built on another germline scaffold, 20 libraries built on other maturated scaffolds, and a completely different target (Fig. 4).

Which physical mechanisms may underly the differences in selective potential that we observe? A number of studies, ranging from structural biology to molecular dynamics simulations, have reported changes in antibody flexibility and target specificity over the course of affinity maturation [28, 29, 30, 31, 32, 33, 34, 35]. The emerging picture is that naïve antibodies are flexible and polyspecific and become more rigid and more specific as they undergo affinity maturation. An increase of structural rigidity in the course of evolution is also found in proteins unrelated to antibodies [36]. Germline scaffolds may thus be more flexible than maturated scaffolds. If this scenario is correct, how this structural flexibility translates into evolutionary diversity once different complementary determining regions (CDRs) are grafted onto the scaffolds remains to be explained.

Irrespectively of mechanisms, we described selectivity distributions of libraries with two statistical parameters, *σ* and *µ*, which we showed to be determined by different factors. Of these two parameters, *σ*, which has simple physical and information-theoretic interpretations, is candidate to serve as a general quantitative index of selective potential for biomolecules. Beyond selection, a next step is to extend this work to quantify evolvability, i.e., the response to successive cycles of selection and mutations. Yet, being able to quantify the selective potential of a scaffold by an index that is systematically reduced in the course of evolution already raises an interesting challenge: can we increase this index to design libraries with better response to selection?

## Acknowledgments

This work was supported by FRM AJE20160635870 and by ANR-17-CE30-0021-02. It benefited from the expertise of the high-throughput sequencing platform at the Institut de Biologie Intégrative de la Cellule (I2BC) at Gif-sur-Yvette, France.

## BOX – Principles of antibody selection experiments

We perform phage display experiments with different libraries of antibodies as input and different molecular targets (DNA hairpins or proteins) as selective pressures [15]. Our antibodies are single domains from the variable part of the heavy chain (V_H_) of natural antibodies. Antibodies in a library share a common scaffold of ≃ 100 amino acids and differ only at four consecutive sites of their third complementary determining region (CDR3), which is known to be important for binding affinity and specificity. A library comprises all combinations of amino acids at these four sites and therefore consists of a total of *N* = 20^4^ ≃ 10^5^ distinct sequences *x* = (*x*_1_, *x*_2_, *x*_3_, *x*_4_). Initial populations include a total of 10^11^ sequences, corresponding to *∼* 10^6^ copies of each of the distinct *∼* 10^5^ sequences when a single library is considered. Physically, these populations are made of phages, each presenting at its surface one antibody and containing the corresponding sequence.

An experiment consists in a succession of cycles, each composed of two steps. In the first step, the phages are in solution with the targets, which are attached to magnetic beads and in excess relative to the phages to limit competitive binding (see Appendix 1.1). The beads are retrieved with a magnet and washed to retain the bound antibodies. In the second step, the selected phages are put in presence of bacteria which they infect to make new phages, thus amplifying retained sequences. A population of *∼* 10^11^ phages is thus reconstituted. Both the selection for binding to the target and the amplification can possibly depend on the sequence of the antibody.

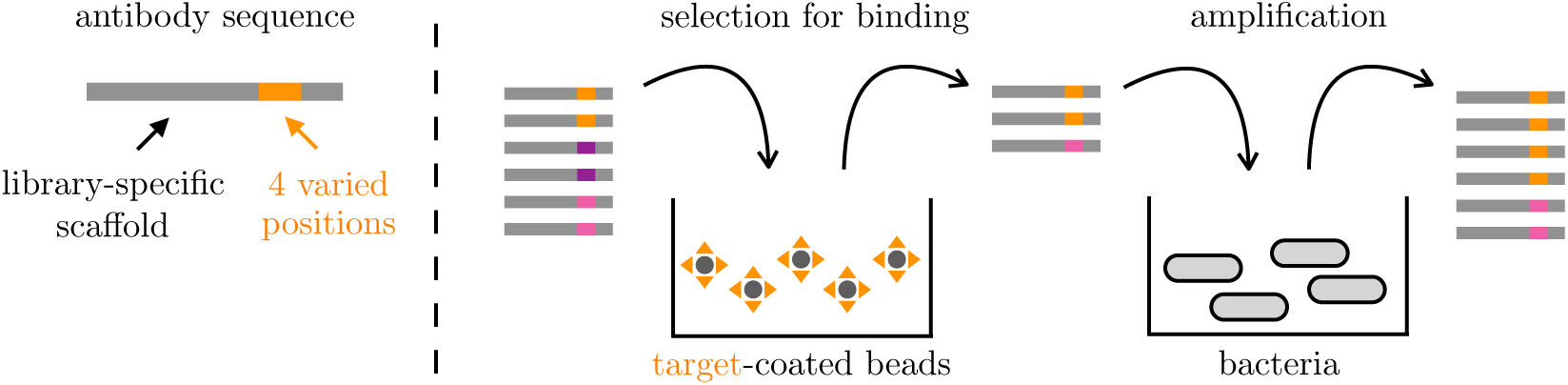

We define the selectivity *s*(*x*) of sequence *x* to be proportional to the probability for sequence *x* to pass one cycle. As the targets are in excess relative to the antibodies, selectivities are independent of the cycle *c* (see Appendix 1.1). In the limit of infinite population sizes, *s*(*x*) is proportional to the relative enrichment *f*^*c*^(*x*)*/f*^*c*−1^(*x*) of the frequencies *f*^*c*^(*x*) after any two successive cycles *c* − 1 and *c*. To estimate these selectivities, about 10^6^ sequences are sampled before and after a cycle and read by high-throughput sequencing. Given the counts *n*^*c*−1^(*x*) and *n*^*c*^(*x*) of sequence *x* before and after cycle *c*, we estimate the selectivity of *x* as

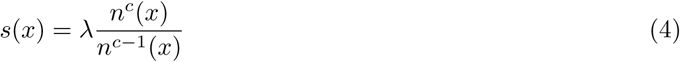

where *λ* is an arbitrary multiplicative factor.

In practice, two types of noise must be taken into account when applying Eq. (4): an experimental noise, which implies that antibodies have a finite probability to pass a round of selection independently of their sequence, and a sampling noise, which arises from the limited number of sequence reads. This sampling noise is negligible if *n*^*c*−1^(*x*) and *n*^*c*^(*x*) are sufficiently large. This is generally not the case for any sequence at the first cycle *c* = 1 where all *N* = 20^4^ sequences are present in too small numbers but becomes the case at the third cycle *c* = 3 for the 100 to 1000 sequences with largest selectivities. We therefore compute *s*(*x*) between the second and third cycles as *s*(*x*) = *λn*^3^(*x*)*/n*^2^(*x*) by restricting to sequences *x* that satisfy *n*^2^(*x*) ≥ 10 and *n*^3^(*x*) ≥ 10. Additionally, as the smallest selectivities are due to experimental noise, we retain only the sequences with *s*(*x*) *> s*\* where *s*\* is determined self-consistently (Appendix 3.2 and Fig. S3). Selectivities *s*(*x*) obtained by this procedure generally depend on the library (scaffold) *L* and the target *T* but are reproducible between independent experiments using the same library and the same target (Fig. S4).

To visualize the sequence dependence of selectivities, we use sequence logos [37]. In this representation, for each position *i* along the sequence, a bar of total height 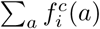 ln 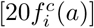 is divided into letters, where each letter represents one of the 20 amino acids *a* with a size proportional to 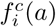, the frequency of *a* at position *i* in the population after cycle *c*; for instance,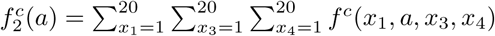; finally, the letters are colored by chemical properties: polar in green, neutral in purple, basic in blue, acidic in red and hydrophobic in black. It illustrates how some motifs are progressively enriched over successions of selective cycles. This representation is, however, dependent on the frequencies *f*^0^(*x*) of sequences in the initial population. To eliminate this dependency, we define an effective frequency 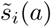 per position *i* and amino acid *a* as 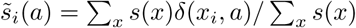, which would correspond to the frequency of *a* at position *i* after one round of selection if all sequences *x* were uniformly distributed in the initial population. It can also be represented by a sequence logo but depends only on *s*(*x*), as illustrated here by the Germ library selected against the DNA1 target (see Figs. S5-S7 for other cases):

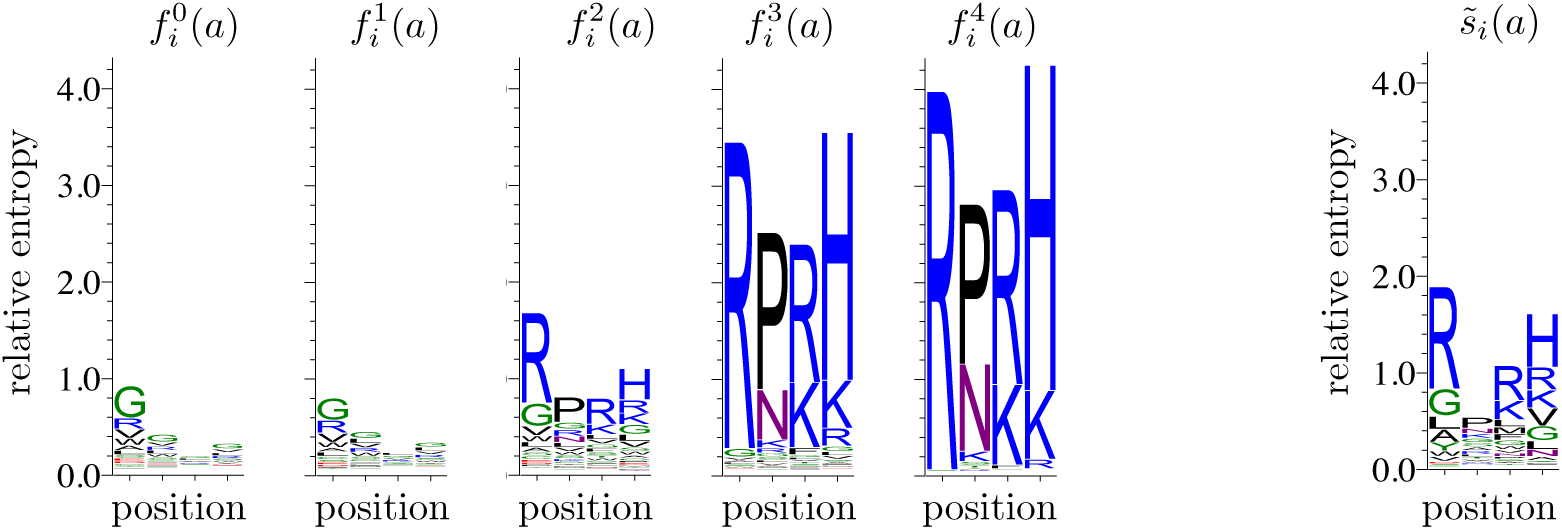

## SUPPLEMENTARY INFORMATION

### 1. Theoretical methods

#### 1.1. Physics of selection

##### 1.1.1. Selectivities and binding energies

When assuming that selection is controlled by equilibrium binding to the target, the distribution of selectivities is constrained by physical principles. Starting with a population of identical antibodies *A* and a single target *T* in excess relative to antibodies, [*T*]_tot_ ≫ [*A*]_tot_, the probability for an antibody to be bound to a target is

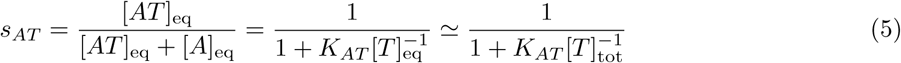

where [*AT*]_eq_ and [*A*]_eq_ are, respectively, the equilibrium concentration of bound and free antibodies and where *K*_*AT*_ = [*A*]_eq_[*T*]_eq_*/*[*AT*]_eq_ is the dissociation constant that characterizes the equilibrium. We used here the fact that most of the targets are unbound so that [*T*]_eq_ = [*T*]_tot_ − [*AT*]_eq_ ≃ [*T*]_tot_, which is justified for our experiments where the total number of targets far exceeds the total number of antibodies, [*AT*]_eq_ *<* [*A*]_tot_ ≪ [*T*]_tot_. The dissociation constant can also be written as *K*_*AT*_ = *k*_−_*/k*_+_, where *k*_+_ and *k*_−_ denote respectively the association and dissociation rates of an antibody-target pair.

We can equivalently write

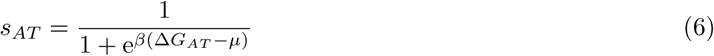

by introducing a binding free energy Δ*G*_*AT*_ = *β*^−1^ ln *K*_*AT*_ and a chemical potential *µ* = *β*^−1^ ln[*T*]_tot_, where *β* sets the energy scale [38]. This Fermi-Dirac statistics is approximated by a Boltzmann statistics

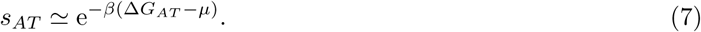

when Δ*G*_*AT*_ ≫ *µ*. This approximation is justified when [*T*]_tot_ ≪ *K*_*AT*_ or, equivalently, [*AT*]_eq_ ≪ [*A*]_eq_, i.e., when the concentration of the targets or the binding affinity are sufficiently low for most of the antibodies to be unbound. Working in this regime is important for the selectivities to reflect binding free energies. Otherwise, the targets are saturating, which cause antibodies to be bound with high probability irrespectively of their dissociation constant.

These conclusions are unchanged when considering a population consisting of different antibodies *A* with different dissociation contants *K*_*AT*_ and binding free energies Δ*G*_*AT*_ = *β*^−1^ ln *K*_*AT*_. In summary, when considering different antibodies *A*, each with its own dissociation constant *K*_*AT*_, the choice of the target concentration [*T*]_tot_ is subject to the two constraints

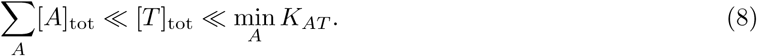

The first constraint Σ*A*[*A*]_tot_ ≪ [*T*]_tot_ guarantees an absence of competition between antibodies so that the selectivities *s*_*AT*_ are intrinsic properties of the sequences of *A*, independent of the composition of the population and therefore independent of the round *c* when successive cycles of selection are performed; formally, [*T*]_eq_, which depends on all *A* present, can then be replaced by [*T*]_tot_ in Eq. (5). The second constraint [*T*]_tot_ ≪ min_*A*_ *K*_*AT*_ guarantees that even the best binders are not in a saturation regime with *s*_*A*_ ≃ 1 independently of differences in their dissociation constants *K*_*AT*_. In our phage display experiments, Σ*A*[*A*]_tot_ ≃ 10^11^ mL^−1^ and [*T*]_tot_ ≃ 10^14^ mL^−1^, which satisfies the first constraint. The concentration Σ*A*[*AT*]_eq_ of selected antibodies before amplification is estimated between 10^5^ mL^−1^ at the first round of selection and 10^7^ − 10^8^ mL^−1^ at the fourth. Considering this last number to reflect properties of the best binders, we estimate that min_*A*_ *K*_*AT*_ */*[*T*]_tot_ ≃ Σ_*A*_[*AT*]_eq_*/* Σ*A*[*A*]_tot_ ≃ 10^3^, which satisfies the second constraint.

##### 1.1.2. Justification and limitations of log-normal distributions

Assuming an additive model for the interaction where the binding energy between sequence *x* = (*x*_1_, *…*, *x*_*𝓁*_) and its target takes is of the form 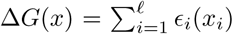 with the *ϵ*_*i*_(*x*_*i*_) taking random values, the central limit theorem indicates that for sufficiently large *𝓁* the energies Δ*G*(*x*) are distributed normally with a mean *µ* ≃ −*𝓁*⟨*ϵ*⟩ and a variance *σ*^2^ ≃ *𝓁*(⟨*ϵ*^2^⟩ −⟨*ϵ*⟩_2_), where ⟨*ϵ*⟩ and ⟨*ϵ*^2^⟩ −⟨*ϵ*⟩_2_ are respectively the mean and variance of the values of binding energies per position *ϵ*_*i*_(*x*_*i*_). Given Eq. (7), this leads to a log-normal distribution for the selectivities *s*(*x*) ∝ *e*^−*β*Δ*G*(*x*)^.

The assumptions involved in this derivation may not be justified, starting from the assumption that selectivity can be equated to binding affinity. However, essentially all deviations from this model, sequence-dependent amplification differences, saturation of the targets, multiple binding sites or non-additive interactions, can be incorporated in a more refined model, at the expense of introducing additional parameters [39]. Deviations from a log-normal distribution of selectivities can therefore, at least in principle, be systematically analyzed and understood.

#### 1.2. Statistics of selection

##### 1.2.1. Extreme value statistics

Extreme value theory states that for any probability distribution P(*S*), the probability to have *S* = *s* ≥ *s*\* conditioned to *S* ≥ *s*\* converges to a generalized Pareto distribution *f*_*κ,s*_*\*,τ* (*s*) = *τ* ^−1^*f*_*κ*_ ((*s* − *s*\*)*/τ*) as *s*\* → ∞ [40], where

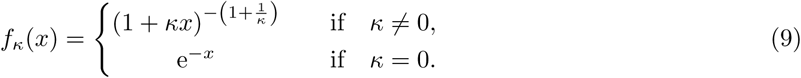

The shape parameter *κ* is determined by the tail of the distribution of *S*. In particular, *κ <* 0 for bounded distributions and *κ* = 0 for distributions with exponentially decreasing tails, including log-normal distributions. On the other hand, *κ >* 0 for distributions whose tail decays as a power-law. For such distributions, when considering a large number *N* of random values *s*_1_ *> s*_2_ *> … > s*_*N*_, *s*_*r*_ *∼ s*_1_*r*^−*κ*^ for *r* ≪ *N*, which is represented in a log-log plot of *s*_*r*_ versus the rank *r* by the linear relationship ln(*s*_*r*_*/s*_1_) *∼* −*κ* ln *r* for the smallest values of *r*.

##### 1.2.2. Effective shape parameter of log-normal distributions

In the asymptotic limit where *N* → ∞ followed by *s*\* → ∞, log-normal distributions are described by a shape parameter *κ* = 0, but their tail decays only slowly. As a result, a large but finite number *N* of random values drawn from a log-normal distribution may appear to be drawn from a distribution with a non-zero shape parameter *κ*_*N*_ ≠ 0.

More precisely, it can be shown that *N* values *s*_1_ *> s*_2_ *> … > s*_*N*_ drawn from a log-normal distribution with parameters *σ, µ* satisfy for *r* ≪ *N* the relation

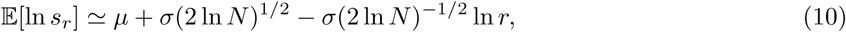

which corresponds to an apparent shape parameter *κ*_*N*_ = *σ*(2 ln *N*)^−1*/*2^ [26]. As *κ*_*N*_ vanishes only very slowly with *N*, it is difficult to determine whether *N* data points arise from a log-normal distribution or from a distribution with a shape parameter *κ >* 0. For instance, increasing the sample size from *N* = 10^5^ to *N* = 10^6^ changes *κ*_*N*_ by only 8 %.

Eq. (10) itself assumes that *N* is large enough. Numerically, we observe that for a given value of *N*, it breaks down when *σ* is below some value *σ*\*. In such cases, the data may appear to arise from a bounded distribution with *κ*_*N*_ *<* 0. Fig. S14 shows the relationship between *κ*_*N*_ and *σ* obtained from numerical simulations when fixing *N* = 10^4^ and *µ* = 0, in which case *σ*\* ≃ 0.5. The same relationship appears as a black dotted line in Fig. 4.

#### 1.3. Information theory of selection

##### 1.3.1. Relative entropies

A general statistical approach to quantify how random variables drawn from a probability *P* ^1^ are consistent with a reference probability distribution *P*^0^ is to use their relative entropy *D*(*P* ^1^‖ *P* ^0^), also known as the Kullback-Leibler divergence [41], which is defined by

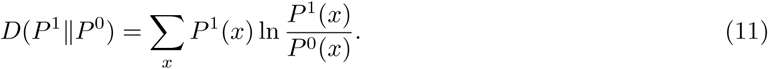

The inverse of this quantity corresponds roughly to the number of samples required to discriminate *P* ^1^ from *P* ^0^. More precisely, the probability under *P*^0^ of *N* samples drawn from *P* ^1^ scales as *e*^−*ND*(*P*^ 1‖ *IP* 0) [41].

##### 1.3.2. Information theory of specific interactions

The problem of quantifying specificity arises when two classes of objects or properties *A* and *T* may be associated. If this association is described by the probability *P*^1^(*A, T*) that *A* is associated with *T*, a natural measure of specificity is *D*(*P*^1^‖*P*^0^) where *P*^0^(*A, T*) represents the expectation from random associations. If *P*^0^(*A, T*) = *P*^1^(*A*)*P*^1^(*T*) where *P*^1^(*A*) = Σ*T P*^1^(*A, T*) and *P*^1^(*T*) = Σ*A P*^1^(*A, T*) are the marginal distributions of *A* and *T*, *D*(*P*^1^‖*P* ^0^) corresponds to the mutual information *I*(*A*; *T*) between the ever, generally does not reflect the expectation from random associations and the relevant measure of specificity is therefore generically not captured by a mutual information but by the more general relative entropy *D*(*P*^1^‖*P*^0^).

In the case of association between a set of ligands *A* and a set of targets *T* controlled by equilibrium binding, the probability *P* ^1^(*A, T*) to find *A* bound to *T* is

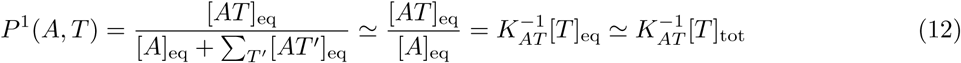

where *K*_*AT*_ is the dissociation constant between *A* and *T* and where the approximations are justified in Appendix 1.1. A random association is defined here by considering equal dissociation constants,

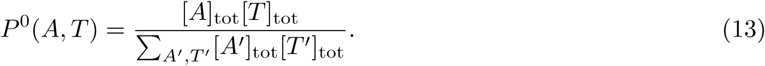

This distribution generally differs from *P* ^1^(*A*)*P* ^1^(*T*).

A selectivity *s*_*AT*_ can be defined for each pair *A, T* as *s*_*AT*_ = *P*^1^(*A, T*)*/P*^0^(*A, T*) so that

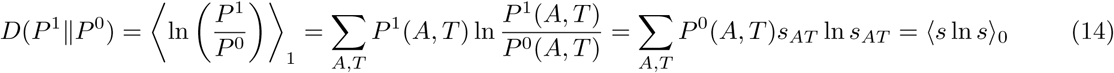

where ⟨·⟩_0_ and ⟨ ·⟩_1_ denote averages taken with *P* ^0^(*A, T*) and *P* ^1^(*A, T*) respectively.

More generally, *s*_*AT*_ = *λP*^1^(*A, T*)*/P*^0^(*A, T*) with an arbitrary multiplicative constant *λ* that can always be written *λ* = ⟨*s*⟩_0_. This corresponds to replacing *s* by *s/*⟨*s*⟩_0_ in the previous formula,

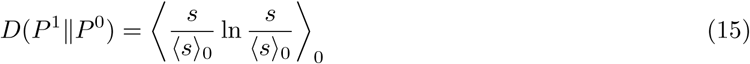

This is equivalent to Eq. (2) where a single target *T* is considered and where *P* ^0^(*A, T*) = 1*/N* and *P* ^1^(*A, T*) = *s*(*x*) with *x* representing the sequence of *A*. This relationship is valid for any initial distribution *f* ^0^(*x*) as long as *f* ^1^(*x*) ∝ *s*(*x*)*f* ^0^(*x*).

The identity

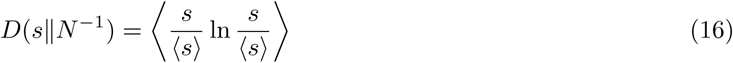

where averages ⟨ ·⟩ are now taken with a distribution *P* (*s*) of the selectivities over the different sequences *x* is, however, valid only when considering as initial distribution a uniform distribution over the sequences. The notation *D*(*s N* ^−1^) assumes, besides, that Σ*x s*(*x*) = 1 so that *s*(*x*) can be interpreted as a probability distribution.

##### 1.3.3. Equivalence with the parameter σ

If further assuming that *P* (*s*) is a log-normal distribution with parameters *σ* and 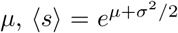 and ⟨*s* ln *s*⟩ = ⟨*s*⟩(*µ* + *σ*^2^) so that

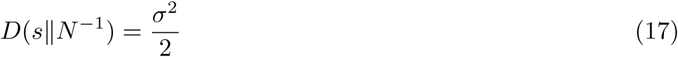

irrespectively of the value of *µ*. This reflects the fact that specificity quantifies only relative differences in binding free energies between different ligands.

A previous study proposed the mutual information as a measure of specificity [27]. It is justified, however, only within the special model considered in [27] where, because of the overall symmetry of the interactions between the *M* locks *A* and *M* keys *T*, *P* ^1^(*A*) ≃ *P* ^1^(*T*) ≃ 1*/M*, and therefore *P* ^0^(*A, T*) = 1*/M* ^2^ ≃ *P* ^1^(*A*)*P* ^1^(*T*).

#### 1.4. Dynamics of selection

##### 1.4.1. Recursion for the sequence frequencies

If *N*^*c*^(*x*) denotes the number of copies of sequence *x* at cycle *c*, the dynamics of selection satisfies the recursion

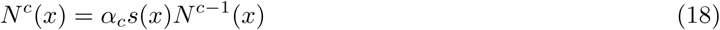

where *α*_*c*_ represents an amplification factor to reach at every round the same total population size *N*, i.e., Σ*x N*^*c*^(*x*) = *N* independent of *c*. In terms of frequencies *f*^*c*^(*x*) = *N*^*c*^(*x*)*/N*, this gives *α*_*c*_ = (Σ*x s*(*x*)*f*^*c*^(*x*))^−1^ and

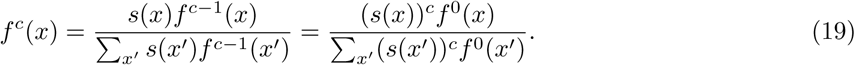

These recursions assume a large *N*, so that the frequencies *f*^*c*^(*x*) = *N*^*c*^(*x*)*/N* are meaningful; in particular, they assume that no sequence disappears.

Note the similarity with a Boltzmann distribution with the cycle *c* playing the role of an inverse temperature.

##### 1.4.2. Recursion for the library frequencies

When considering a population consisting of an equal mix of different libraries *L*, the frequency *f*^*c*^(*L*) = Σ*x*∈*L f*^*c*^(*x*) of library *L* satisfies the recursion

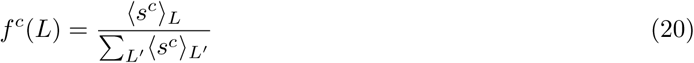

With

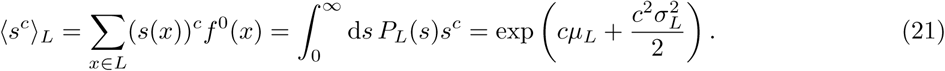

Here, the first equality defines the average ⟨ ·⟩_*L*_ within each library *L*. The second equality, on the other hand, makes two assumptions: first, that selectivities *s* within library *L* are described by a distribution of selectivities *P*_*S*_ (*s*) and, second, that sequences within a library are uniformly represented in the initial population. The third equality makes the additional assumption that *P*_*L*_(*s*) is a log-normal distribution with parameters *σ*_*L*_ and *µ*_*L*_.

Under these different assumptions, the frequency of library *L* at cycle *c* is given by

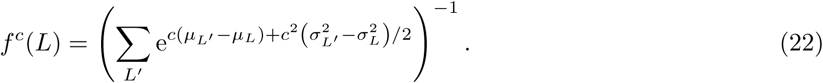

This shows that for small *c*, the dynamics is controlled by the *µ*_*L*_, with in limit *c* → 0, (*f*^*c*^(*L*)−*f*^0^(*L*))*/f*^0^(*L*) ≃ *c*(*µ*_*L*_ − ⟨*µ*⟩), i.e., at the first cycle, the frequency of library *L* increases if its *µ*_*S*_ exceeds the average ⟨*µ*⟩ across libraries and it decreases otherwise. For large *c*, on the other hand, the dynamics is controlled by the *σ*_*L*_s with *f*^*c*^(*L*) → 1 for the library *L* that has largest *σ*_*L*_, regardless of the values of *µ*_*L*_.

These calculations rely on several assumption, in particular the assumption that sequences within a library have initially uniform frequencies, which is not satisfied in the experiments. This explains the differences between the model and the data in Fig. 3.

### 2. Experimental methods

Experimental methods are as in our previous work [14], except for target immobilization and sequencing data analysis as summarized below.

#### 2.1 Phage production

Production of antibody-displaying phage was performed through infection of library cells (TG1 strain) with M13KO7 helper phage and growth at 30°C for 7 h in selective 2xYT medium containing 100 *µ*g*/*mL ampicillin (Sigma-Aldrich, Saint-Louis, MO, USA). Cells were then centrifuged and the supernatant containing displaying phages was kept and stored at 4°C overnight. All selections were performed on the day immediately following the phage production step.

#### 2.2 Target immobilization

Target molecules were immobilized on streptavidin-coated magnetic beads (Dynabeads(R) M-280 Streptavidin) purchased from Invitrogen Life Technologies (Carlsbad, CA, USA). The DNA hairpin targets (DNA1 and DNA2) in fusion with a biotin at their 5′ end were purchased from IDT (Leuven, Belgium) diluted in MilliQ water and stored at −20°C. The genes of protein targets (eGFP and mCherry, corresponding respectively to PDB IDs 2Y0G and 2H5Q) in fusion with a SBP tag were kindly provided by Sandrine Moutel (Institut Curie, Paris, France). They were produced in liquid T7 Express *E. Coli* cultures induced at OD_600_ = 0.5 with 300 *µ*M Isopropyl *β*-D-1-thiogalactopyranoside (IPTG, Sigma-Aldrich, Saint-Louis, MO, USA) final and incubated overnight at 30°C. The proteins were harvested by threefold flash freezing in liquid nitrogen and quick thawing in a water bath at 42°C, followed by incubation with 50 *µ*g*/*mL lyzozyme final and 2.5 U*/*mL DNase I final at 30°C for 15 minutes and centrifugation at 15, 000 g and 4°C for 30 minutes. The supernatant was aliquoted in protein low-bind tubes (Protein LoBind, Eppendorf, Hamburg, Germany), flash frozen in liquid nitrogen and stored at *-*80°C until use.

Binding of target molecules to streptavidin-coated magnetic beads was performed in DNA low-bind tubes (DNA LoBind tubes, Eppendorf, Hamburg, Germany) for the DNA targets or protein low-bind tubes (Protein LoBind tubes, Eppendorf, Hamburg, Germany) for the protein targets. Beads and targets were incubated in 0.5x PBS at ambient temperature on a rocker for 15 min, followed by removal of all liquid and 3 washing steps: addition of 500 *µ*L washing solution, vortexing, separation of beads using a magnet and removal of all liquid. Finally, the beads were stored in washing buffer at 4°C for use on the following day. Bw1X buffer (1 M NaCl, 5 mM Trizma at pH = 7.4, 0.5 mM EDTA) was used as washing buffer for DNA targets (to screen electrostatic interactions), 1x PBS with 0.1 % Tween20 for protein targets (to screen hydrophobic interactions). The same procedure was followed for negative/null selection tubes, with MilliQ water instead of target solutions.

Successful immobilization of protein targets was confirmed by fluorescence measurements of treated beads against untreated and MilliQ water-treated beads as negative controls.

#### 2.3 Phage display selection

The selection protocol is as previously published in [14]. The washing buffer was removed from the target-covered beads. Then, 1 mL of culture supernatant from the phage production step containing ≈ 10^11^ phages was added to the negative selection tube (containing no targets) and incubated for 90 minutes at ambient temperature, shaking. The beads were separated by a magnet and the liquid was transferred to the positive selection tube (containing the targets) and incubated for 90 minutes at room temperature, shaking. Finally, all liquid containing unbound phage was removed and the beads were subjected to a 10-fold washing using 10 mL of 1x PBS with 0.1 % Tween20. Bound phage were eluted from beads with 1.4 % triethylamine (Sigma-Aldrich, Saint-Louis, MO, USA) in MilliQ water and used for infection of fresh exponential TG1 cells to obtain the selected library.

#### 2.4 Illumina sequencing

Glycerol stocks of library cells at relevant selection cycles were defrosted and plasmids were extracted using purification kits from Macherey-Nagel (Düren, Germany). No liquid culture was performed prior to plasmid extraction to avoid potential additional biases from growing an overnight culture beforehand. Resulting plasmids were used as input for Illumina sequencing preparation PCR: a first reaction using primer sequences common to all three libraries downstream CDR_3_ (GCTCGAGACGGTAACCAGG, forward) and halfway inside V_H_ (ACAACCCGTCTCTTAAGTCTCGT, reverse) added random barcodes of length 5 nt to discriminate between neighboring clusters. A second reaction added P5 and P7 indices to identify library, target and selection round corresponding to each cluster, as well as the adapter for the sequencing procedure. Illumina sequencing and demultiplexing were performed at I2BC, Gif-sur-Yvette, France.

### 3. Data analysis

#### 3.1 Preprocessing

The Illumina sequencing yields for each sample (i.e., each library, target and selection round) between 10^5^ and 5.10^6^ sequencing clusters. The data files contain the entirely overlapping forward and reverse reads for all clusters of a given sample. Each cluster was accepted or discarded based on the following procedure: Both the forward and reverse reads were screened for the presence of the primer sequences (up to 4 nt mismatch accepted for each) and cut to keep only the part between the primers (including the primers). Either one was discarded if the primer search was unsuccessful. We then checked if the remaining forward and/or reverse sequence fragments have the expected length of 170 nt, corresponding to the region of interest. If only one direction had the expected length, only this direction was kept and the other one was discarded. If both directions did not have expected length, the complete cluster was discarded. Finally, if both reads had expected length, a consensus sequence was generated by taking on each position with disagreement between both reads the nucleotide measured with highest quality read. A final check was performed for (i) a sufficient average quality read over the whole region of interest (⟨*Q*⟩ ≥ 59) and (ii) the restriction sites immediately up- and downstream CDR_3_ (TGTGCGCGC and TTCGACTAC) are located at their expected positions (108-116 and 129-137 in reverse direction; up to 4 nt mismatch accepted for each). The cluster was discarded if either of these two criteria was not fulfilled.

After completion of this procedure, (i) the framework (Germ, Lim or Bnab) and (ii) the CDR3 sequence for all remaining sequencing reads in the full-library experiments were identified. Step (i) was performed by measuring the Hamming distance of the visible library-specific framework part upstream the CDR3 of the read (of length 116 nt) to all three framework reference sequences. The read was assigned to the nearest framework if the Hamming distance to the nearest framework was ≤ 7 nt *and* the difference in Hamming distance to the nearest and next-nearest frameworks was ≥ 3 nt. For step (ii), the CDR3 sequence was simply extracted from the read for the full-library experiments. For the selections with reduced diversity a similar procedure as for the framework part was applied: the measured CDR3 sequence was assigned to the nearest among ∼ 20 reference sequences if the Hamming distance was ≤ 3 nt and the difference in Hamming distance between nearest and next-nearest was ≥ 1 nt. After assessment of the sequence identity of all clusters in a dataset, the CDR3 sequences were translated into amino acids and the number of occurrences of each clone (determined by its framework and its CDR3 sequence) was counted.

The nucleotide sequences of the visible framework parts upstream the CDR3 of all three libraries as well as the Hamming distances *d*_*H*_ between the pairs is as follows:

Germ:

~~~
ACAACCCGTCTCTTAAGTCTCGTGTTACCATCTCTGTTGACACCTCTAAAAACCAGTT…
CTCTCTGAAACTGTCTTCTGTTACTGCGGCGGACACTGCGGTTTACTACTGTGCGCGC
~~~

Lim:

~~~
ACAACCCGTCTCTTAAGTCTCGTGTTACCATCTCTATCGACACCTCTAAAAACCACTT…
CTCTCTGCGTCTGATCTCTGTTACTGCGGCGGACACTGCGGTTTACCACTGTGCGCGC
~~~

Bnab:

~~~
ACAACCCGTCTCTTAAGTCTCGTCTGACCCTGGCGCTGGACACCCCGAAAAACCTGGT…
TTTCCTGAAACTGAACTCTGTTACTGCGGCGGACACCGCGACCTACTACTGTGCGCGC
~~~

*d*_*H*_ (Germ, Lim) = 10 nt, *d*_*H*_ (Lim, Bnab) = 25 nt and *d*_*H*_ (Germ, Bnab) = 22 nt.

For the mixed full-library selections, final data files contain three columns: 1) framework identity (‘`germ`’ for Germline, ‘`lmtd`’ for Limited, ‘`bnAb`’ for Bnab, ‘`????`’ if framework inference failed), 2) CDR3 identity given by the sequence of 4 amino acids or the sequence of 12 nucleotides or by ‘`????`’ if the CDR3 readout failed, 3) number of occurrences in the dataset. The preprocessed data from the experiments reported in this paper is made available in this format.

We checked that the results are unaffected by the choice of the parameters in the preprocessing procedure described here.

#### 3.2 Noise cleaning

Selectivities are computed from sequencing counts as indicated in Eq. 4. To account for sampling noise, only sequences whose count is ≥ 10 both at round *c* and *c*+1 are considered. Moreover, we ignore selectivities *s*(*x*) below a threshold *s*\*, which arise from unspecific binding. Unspecific binding modifies the expression for the selectivity of sequence *x* to include a sequence-independent unspecific binding energy Δ*G*_us_,

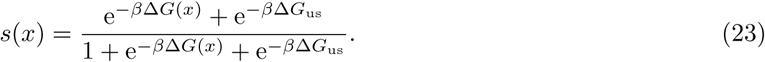

It sets a lower bound for the selectivity given by

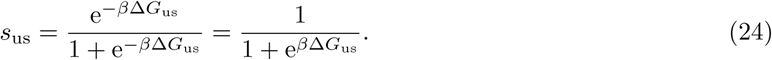

The argument for log-normality of selectivity distributions applies only when the specific binding contribution Δ*G*(*x*) dominates the selectivity. We therefore eliminate the selectivities dominated by unspecific binding.

This is done by introducing a cut-off *s*\*. The choice is made such that (i) the values of the inferred parameters 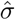 and 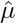 are approximately constant for all *s* ≥ *s*\* and (ii) *s*\* is large enough to eliminate selectivities due to unspecific binding. This last condition is implemented by plotting the counts *n*^2^(*x*) and *n*^3^(*x*) at the two successive cycles, as illustrated in Figure S3: sequences with *s* = *s*_us_ appear in the diagonal with a variance that decreases with increasing counts, as expected from sampling noise, and *s*\* is chosen so as to exclude these sequences. In cases where specific binding to the target is very strong, sequences selected for unspecific binding are not present (Fig. S15A), while in cases where specific binding is too weak, only sequences selected for unspecific binding are present (Fig. S15F).

The same criteria apply when fitting to generalized Pareto distributions to infer the parameter *κ* but criterion (i) may lead to a higher value of *s*\* if the measured selectivities extend beyond the tail of the distribution. In our previous work [14], we only considered criterion (i). In one case (Frog3 against DNA1), the *s*\* that we define here by accounting for (ii) differs from the *s*\* that had previously defined (Fig. S15), which leads to a significantly different estimation of 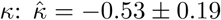 instead of 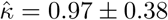. In the other cases, we recover essentially the same results. The new analysis provides, however, additional insights; in the case of Frog3 against PVP, it thus appear that the vanishing value of *κ* can be attributed to the selectivities being dominated by unspecific binding (Fig. S15).

#### 3.3 Fit to log-normal distributions

To infer from experimental data the parameters *σ* and *µ* of a log-normal distribution, as given by Eq. (1), we focus on the best available selectivities *s*_*i*_ *> s*\*, the log-normal distribution is under-sampled. In practice, it is more convenient to work with the log of the selectivities, *y*_*i*_ = ln *s*_*i*_, and to fit them with a normal distribution. If restricting to values *y*_*i*_ larger than a given threshold *y*\*, the probability ℙ [*Y* = *y*|*Y* ≥ *y*\*] of observing *y*_*i*_ given that *y*_*i*_ ≥ *y*\* is

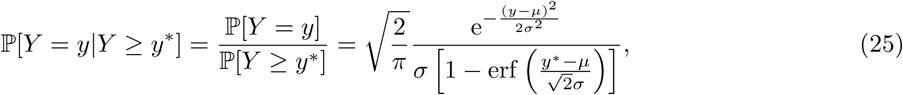

where erf 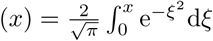 is the Gauss error function. The log-likelihood ℒ (*µ, σ, y*\*) then verifies

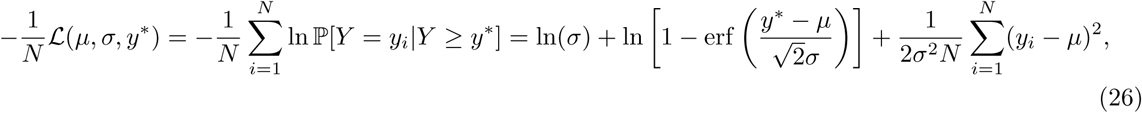

up to irrelevant additive constants independent of the parameters *µ* and *σ*. For a given *y*\*, we minimize this quantity with respect to the parameters *σ* and *µ* to obtain 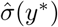 and 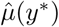 and then chose *y*\* such that for any *y* ≥ *y*\* both 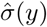 and 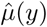 are nearly constant (criterion (i) in Appendix 3.2). Finally, we obtain a lower bound on the uncertainty of the parameter values using the Fisher information matrix and the Cramer-Rao bound. To assess the quality of fit, we produce P-P plots comparing the cumulative distribution of data to

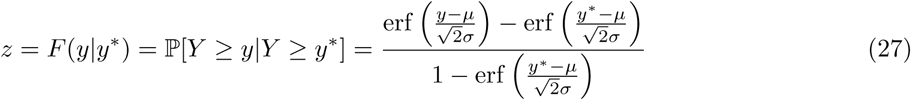

where *z* is the fraction of the data above *y* ≥ *y*\* according to the model, and Q-Q plots comparing the data to the inverse distribution function *y* = *F* ^−1^(*z*|*y*\*).

#### 3.4 Normalization of µ across libraries

The selection of a library *L* against a target *T* yields only the values of the highest selectivities *s*(*x*) up to an unknown multiplicative constant *λ* (see Box). The parameter *σ* = *σ*_*L,T*_ is independent of *λ* but not the parameter *µ* = *µ*_*L,T*_. The relative values of *µ*_*L,T*_ for different libraries *L* selected against the same target *T* are determined by performing selections where the different libraries are mixed in the initial population: this leaves undetermined one overall multiplicative constant per target. Finally, we fix them by setting *µ*_Germ,*T*_ = 0 for each target *T*. In practice, inferring *µ* from the tail of *P* (*s*) is challenging, even more so when different libraries are mixed, as one library often dominates the population after a few cycles. To overcome this limitation, we can separately measure the selectivities of random sequences, which typically belong to the mode of the distribution *P* (*s*), located at *m* = *µ* − *σ*^2^.

For a given target, our approach is thus to first perform 3 cycles of selection with each library, Germ, Lim and Bnab. Using the results from cycles 2 and 3, we estimate as many selectivities *s*_*L,T*_ (*x*) as possible (see Box and Fig. 1A). We then identify 2 to 4 sequences with largest selectivity from each library, which we mix with 2 to 4 random sequences from each library, and perform one round of selection of the mixture of these ∼ 20 sequences. From the results of this experiment, we estimate with high precision the relative selectivities of top and typical sequences from the different libraries (Fig. 1B). We typically find that the random sequences from a same library have a similar selectivity which we use to define the relative modes *m*_*L,T*_ of the 3 libraries. Given these modes *m*_*L,T*_, we then infer from the available values of *s*_*L,T*_ (*x*) the parameter *σ*_*L,T*_ by maximum likelihood, using the relationship 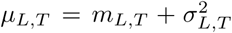. Finally, we fix the remaining overall multiplicative constant by setting *µ*_Germ,*T*_ = 0.

In practice, to reduce the total number of experiments, we performed the selection of the full libraries in mixtures; as we verified with one target, the results are equivalent to those obtained from separate selections (Fig. S8). We also found unecessary to estimate the selectivities of typical sequences against all targets once we understood that these values are not controlled by the target.

## SUPPLEMENTARY FIGURES

**Figure S1:**
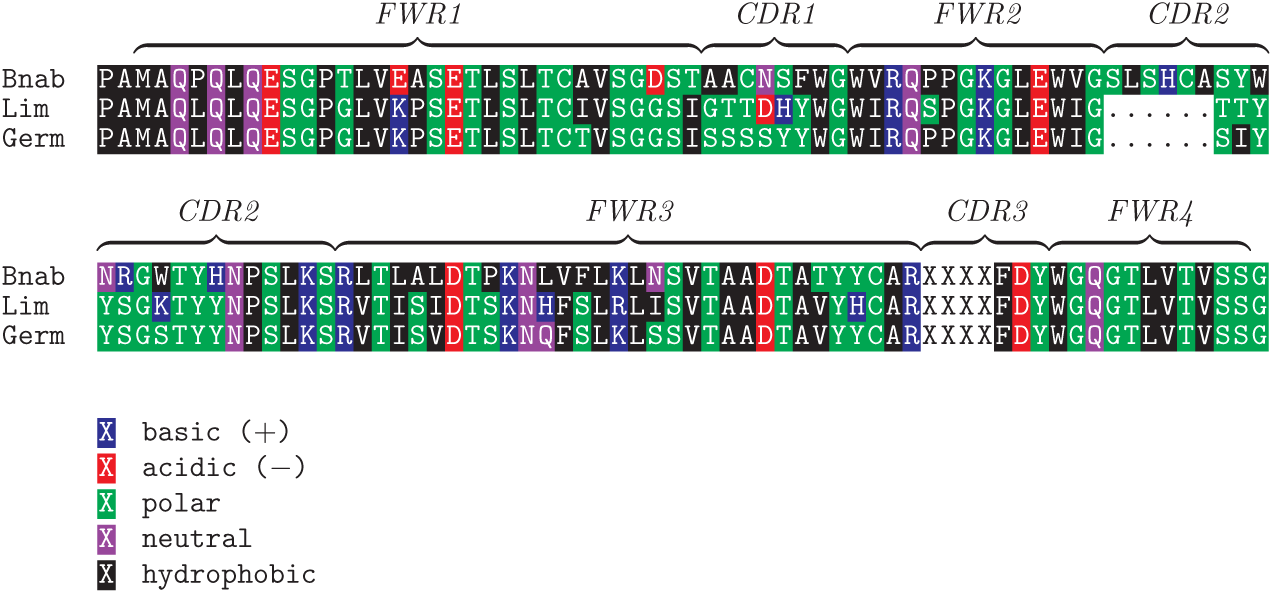
Alignment of the sequences of the three scaffolds, Bnab, Lim and Germ. The 4 randomized positions correspond to the part of the CDR3 indicated by `XXXX`.

**Figure S2:**
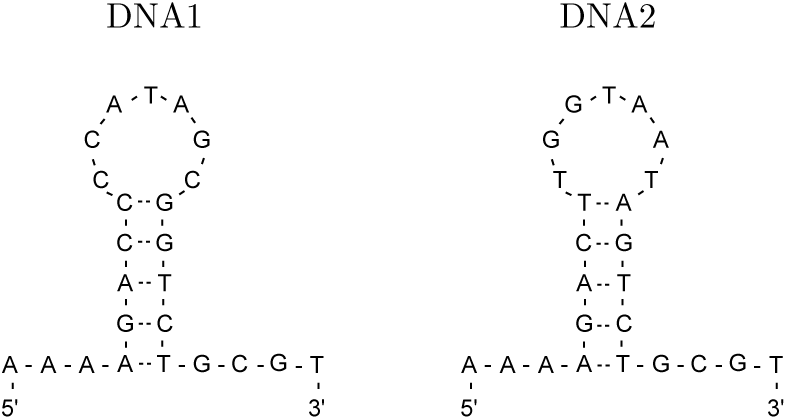
DNA1 and DNA2 binding targets. The targets display a hairpin structure at room temperature. They share a common stem sequence but the sequence of their loop differ. A biotin is placed at the 5’ ends to allow for immobilization on streptavidin-coated magnetic beads.

**Figure S3:**
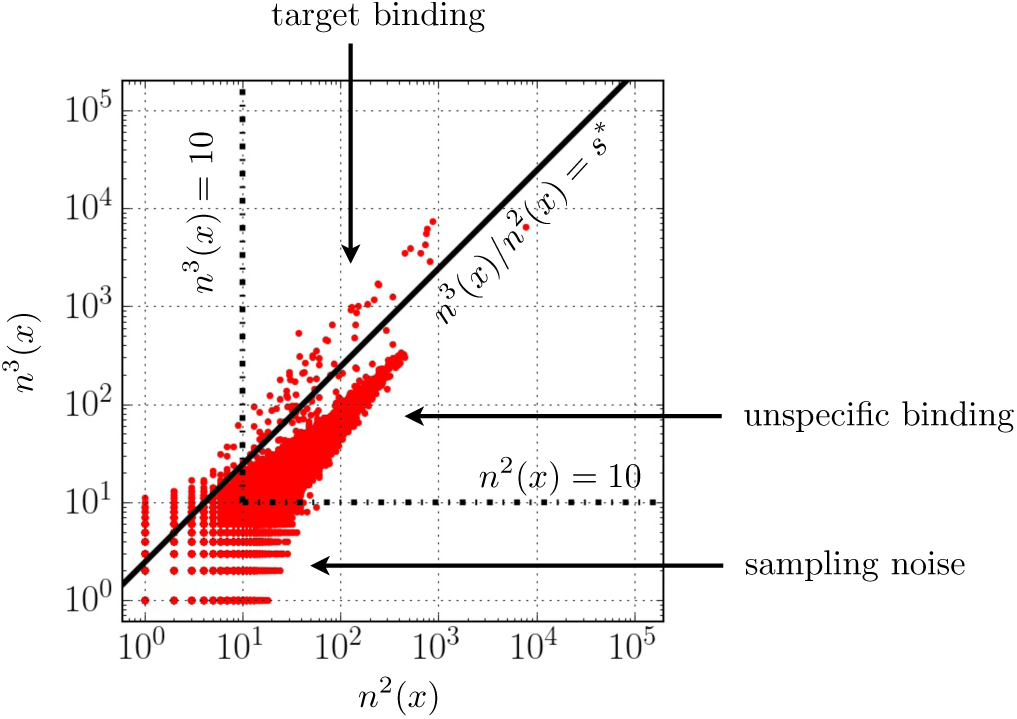
Illustration of the choice of the cutoff *s*\* below which measured selectivities are attributed to unspecific binding. The number *n*^3^(*x*) of counts in the sequencing data at round *c* = 3 is plotted against the number *n*^2^(*x*) of counts at round *c* − 1 = 2 for a selection of the Bnab library mixed with the two other libraries against the DNA1 target. An accumulation of sequences with similar selectivities is observed along the diagonal, with larger variance for smaller values as expected from an increased sampling noise. This is interpreted as arising from unspecific binding, associated with a selectivity *s*_us_ independent of the sequence. We define a cut-off *s*\* such that sequences *x* with *s* = *n*^3^(*x*)*/n*^2^(*x*) ≥ *s*\* cannot be attributed to unspecific binding. In addition, we restrict to sequences *x* with *n*^2^(*x*) ≥ 10 and *n*^3^(*x*) ≥ 10, as represented by the vertical and horizontal lines, to ensure that the inferred selectivities are not dominated by sampling noise.

**Figure S4:**
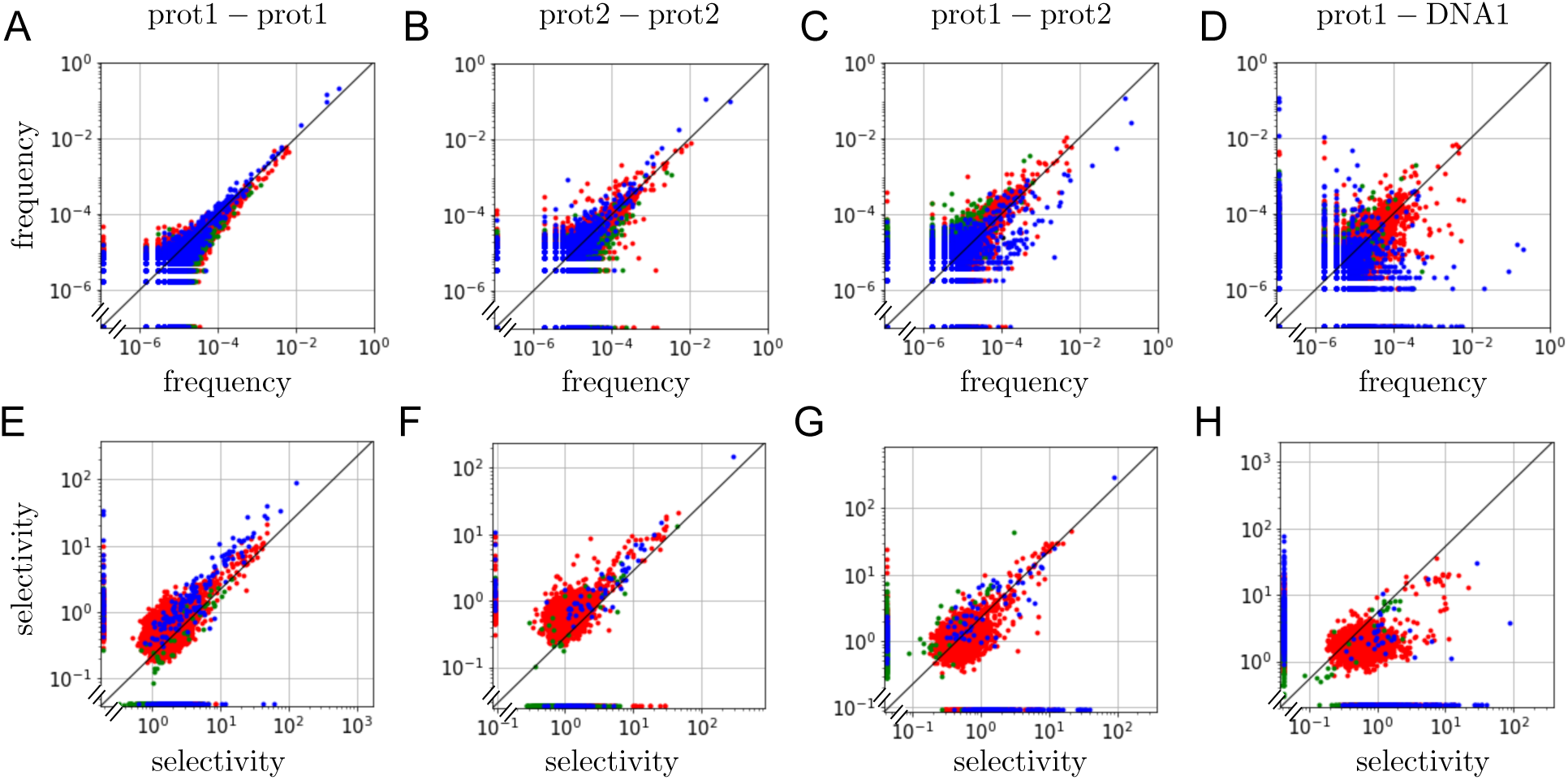
Comparisons between results of replicate and non-replicate experiments. **A.** Comparison of the frequencies *f* ^3^(*x*) = *n*^3^(*x*)*/* _*x*′_ Σ*n*^3^(*x*′) computed after the third cycle (*c* = 3) between two independent replicate experiments where a mixture of the Germ (in blue), Lim (in green) and Bnab (in red) libraries is selected against the protein target prot1. Due to stochastic sampling, some sequences *x* are well represented in one experiment (*n*^3^(*x*) ≥ 10) but not in the other; they are represented by the points along the two axes. As expected, the frequencies of the most prevalent sequences are the most reproducible. **B.** As in A but for protein target prot2. **C.** Comparing an experiment with prot1 as target with another with prot2 as targeϵ common sequences are enriched in the two cases, although with not exactly the same frequencies. **D.** Comparing an experiment with prot1 as target with another with DNA1 as target, showing that different sequences are enriched in each case. In particular, the most frequent sequences when selecting against one target are absent in the third round when selecting against the other (points along the axes). **E**,**F**,**G**,**H.** Comparison of selectivities *s*(*x*) calculated from the frequencies between the second and third rounds as *s*(*x*) = *λn*^3^(*x*)*/n*^2^(*x*). Points along the axes correspond to sequences for which the selectivity could be estimated only for one of the two experiments. We verify that in cases E,F,G where the targets are similar the same top selectivities are recovered (up to a multiplicative constant corresponding to a shift in log-log plots). Beyond stochastic effects, reproducibility is mainly limited by the differences in the production of the targets, as shown in Fig. S12.

**Figure S5:**
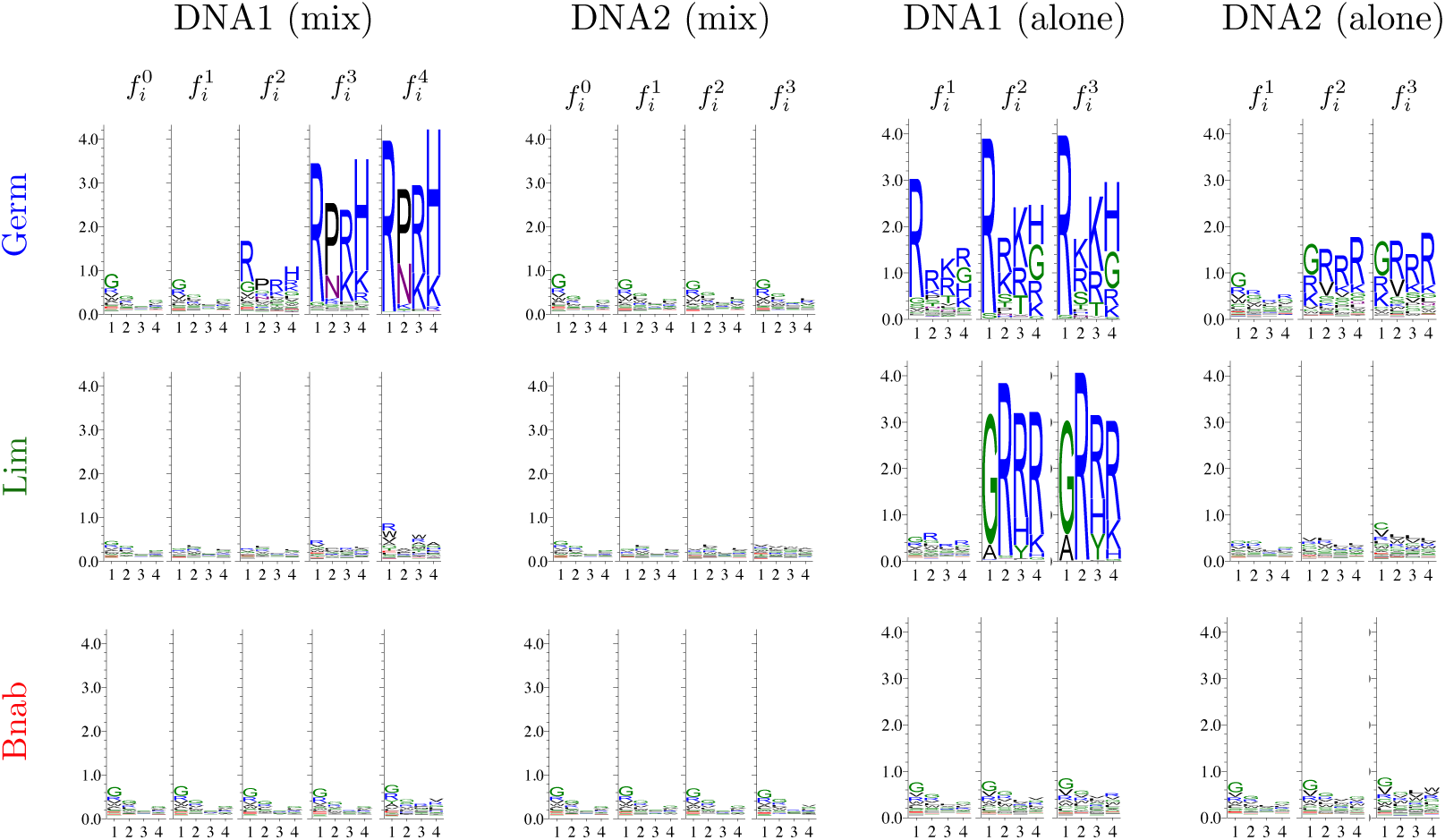
Extension of the figure in the Box to the 3 libraries Germ, Lim, Bnab selected either in a mixture (mix) or on their own (alone) against the DNA1 and DNA2 targets. The sequences logos represent the frequencies 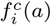 of amino acids at each successive cycle *c* = 0,1,2,3,4.

**Figure S6:**
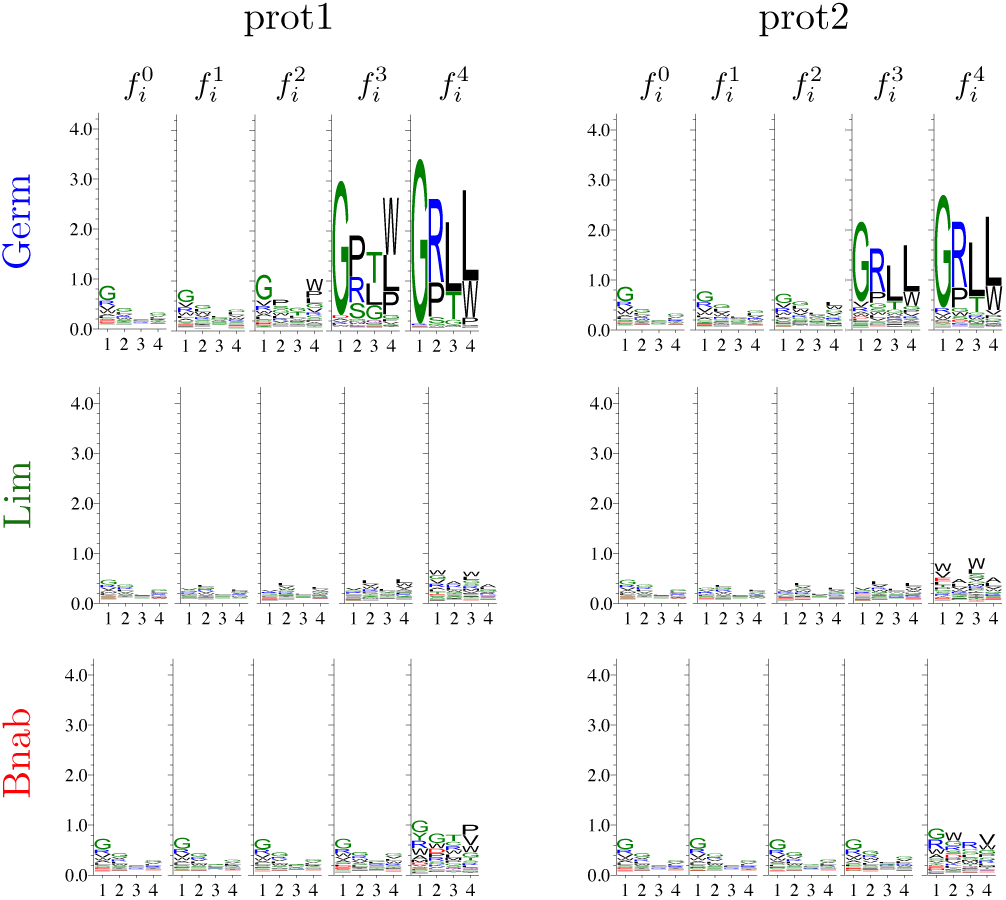
Extension of the figure in the Box to the 3 libraries Germ, Lim, Bnab selected in mixture against the prot1 and prot2 targets. The sequences logos represent the frequencies 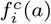 of amino acids at each successive cycle *c* = 0, 1, 2, 3, 4.

**Figure S7:**
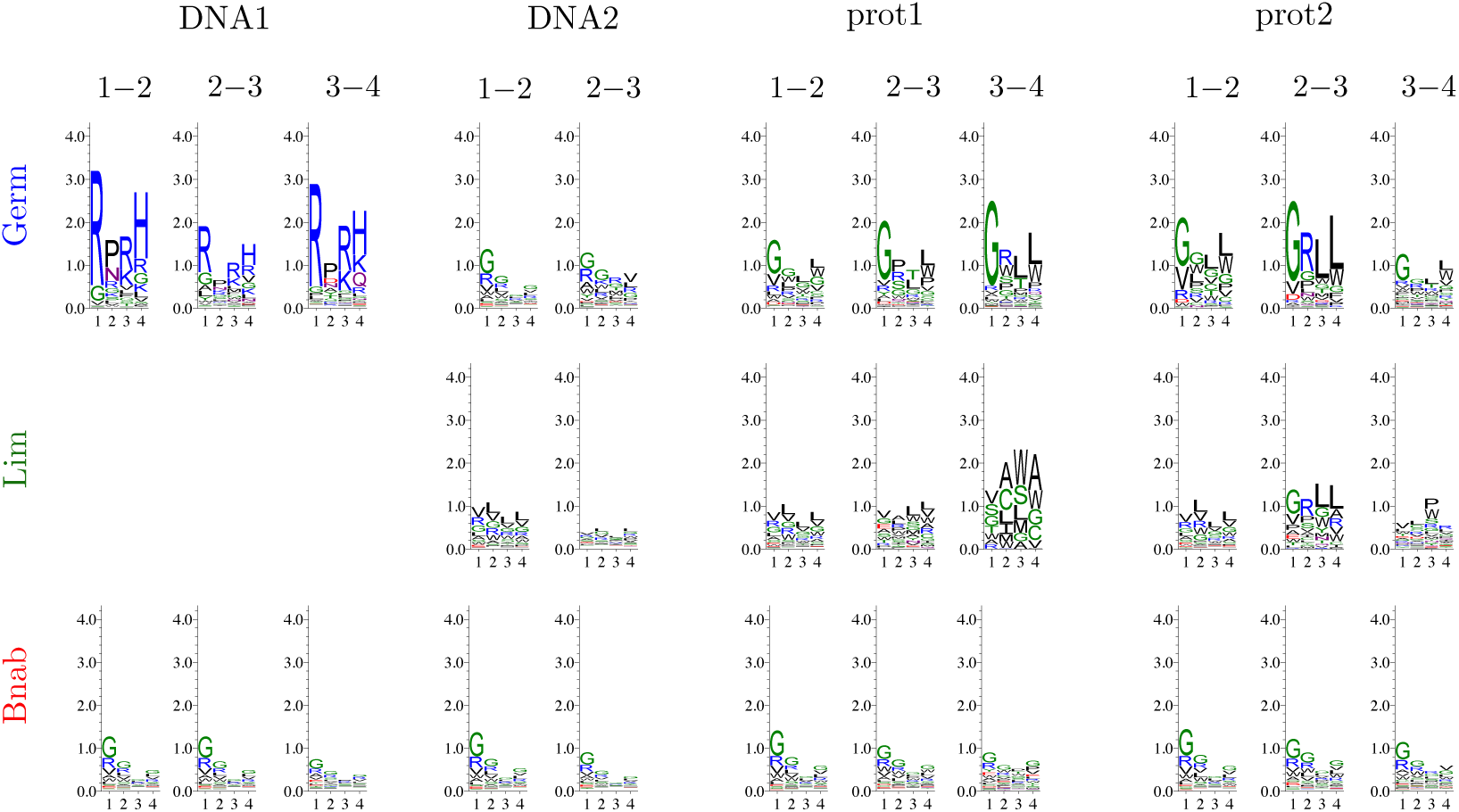
Sequence logos for the selectivities 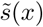 computed between two successive rounds (1-2, 2-3 or 3-4). The differences between rounds reflect sampling fluctuations.

**Figure S8:**
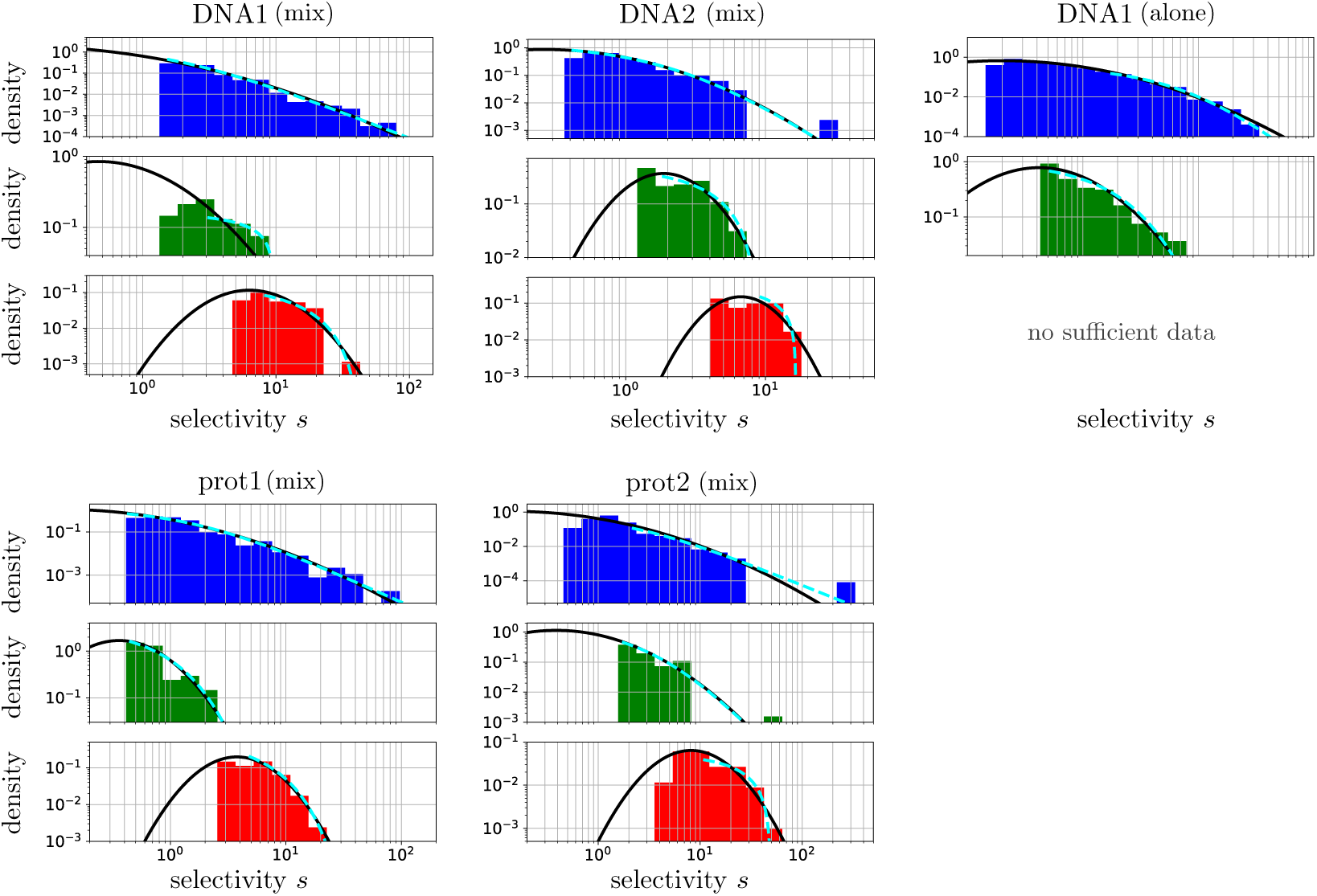
Distributions of selectivities of the three libraries (Germ in blue, Lim in red, Bnab in red) when selected either in a mixture (mix) or on their own (separate) against the different targets. This figure extends Fig. 1A that reports the selection against the DNA1 target of the Germ and Bnab libraries in mixture and of the Lim library on its own. In addition to the best fits to a log-normal distribution (black curves), the best fits to generalized Pareto distributions are also shown (cyan dotted curves). The selection of the Bnab library alone against the DNA1 target yielded insufficient data for a meaningful analysis.

**Figure S9:**
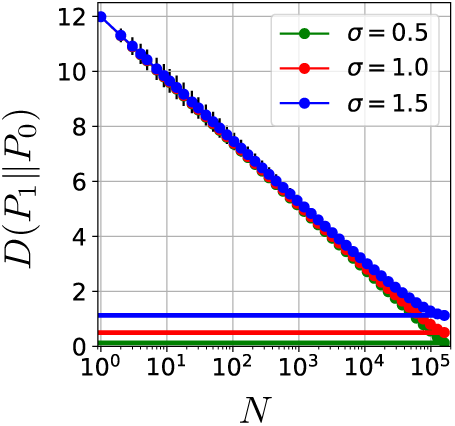
How the estimation of the entropy is biased by finite sampling. 10^5^ values were drawn from a log-normal distribution with parameters *µ* = 0 and *σ* = 0.5 (green), 1 (red) and 1.5 (blue). The relative entropy *D*(*P*_1_ ‖ *P*_0_) was then estimated using a random subsample of size *N*. For any *N <* 10^5^, this leads to an overestimation of *D*(*P*_1_ ‖ *P*_0_) whose actual value *σ*^2^*/*2 (see Eq. (3)) is represented by the horizontal lines at the bottom.

**Figure S10:**
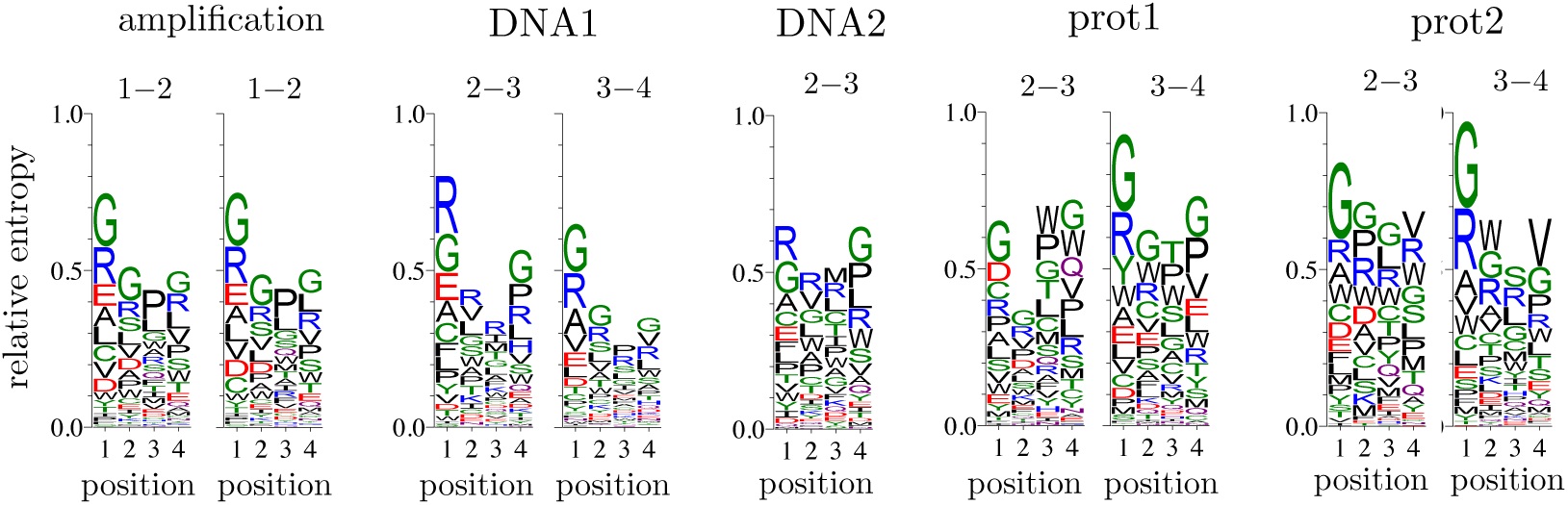
Sequence logos for the selectivities 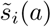 of the Bnab library subject to either amplification only or to amplification and selection for binding against the DNA1, DNA2, prot1 or prot2 targets. The selectivities are computed between the first and second cycles (1-2) or between the third and fourth cycles (3-4); for amplification only, the results of two replicate experiments are shown. The sequence logos of selectivities calculated between rounds 2 and 3 are the same as those shown in Fig. 2 (Bnab library), except for the scale along the y-axis. All sequences logos share common patterns reflecting a common contribution from amplification biases. Sequence logos against the protein targets show, however, an enrichment for tryptophane (symbol `W`) that is not observed when selection involves amplification only. Selections of the Bnab library thus have a target-dependent contribution from binding affinity of similar order of magnitude as a common target-independent contribution from amplification biases.

**Figure S11:**
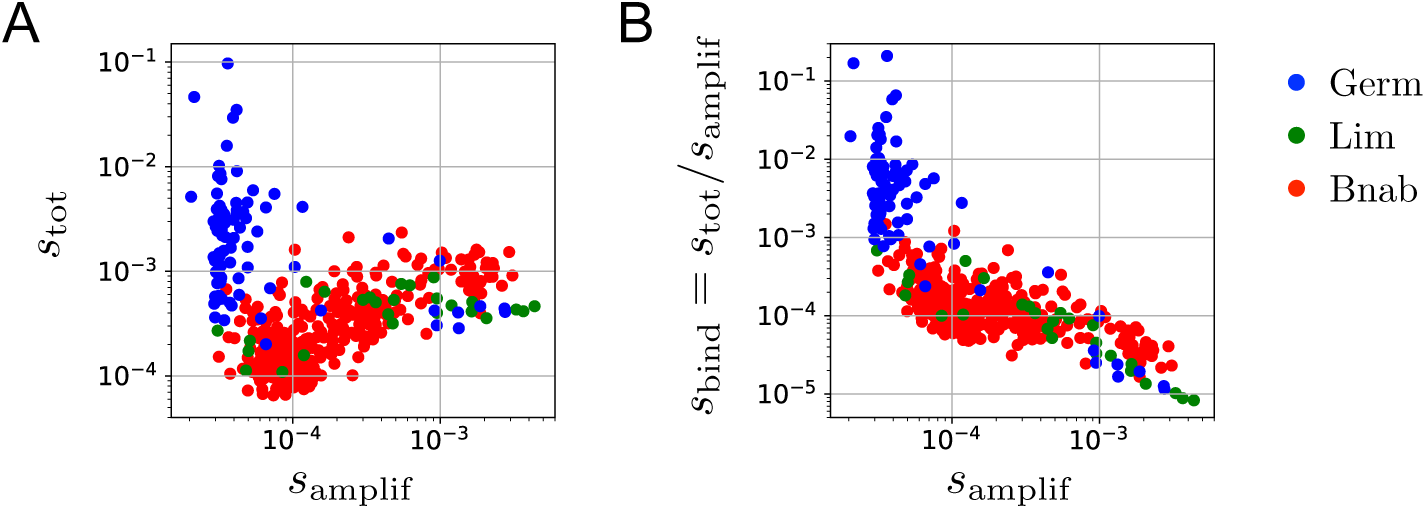
Contribution of amplification biases to the selectivities in selection against the DNA1 target. A separate experiment without any selection for binding was performed to estimate the difference of selectivities arising from the amplification step alone. **A.** The resulting *s*_amplif_ is here compared to the selectivities *s*_tot_ from an experiment including a selection for binding. The sequences with top *s*_tot_, which all belong to the Germ library (in blue), are among the sequences with lowest *s*_amplif_, which indicate that they are selected for binding with no contribution from the amplification bias. On the other hand, the sequences with top *s*_tot_ from the Lim and Bnab libraries (respectively in green and red), have also top *s*_amplif_, which indicate a significant contribution from amplification biases. **B.** The ratio *s*_tot_*/s*_amplif_ represents the contribution to selectivity of binding alone. The two selective pressures, binding and amplification, appear here to be orthogonal.

**Figure S12:**
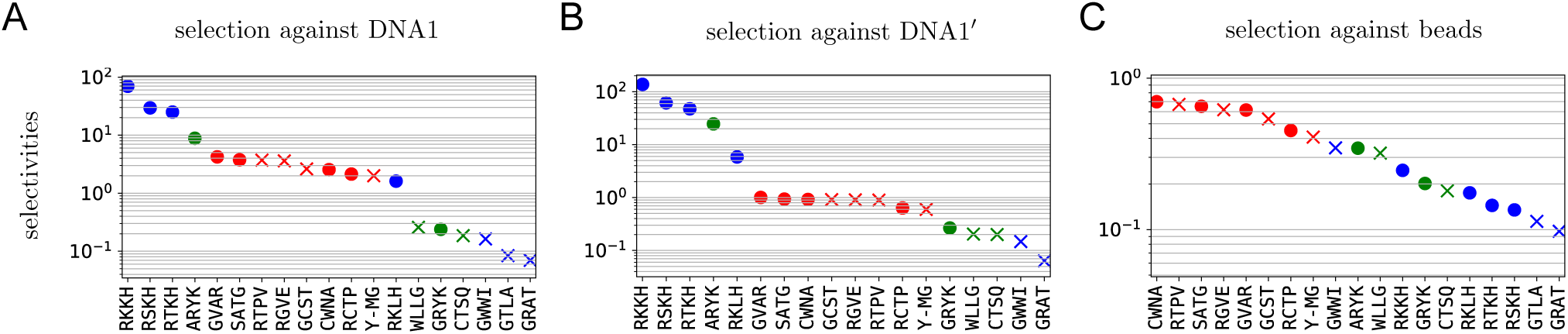
Supplementary experiments with minimal libraries. **A.** Selectivities of top and random sequences from the three libraries, Germ (in blue), Lim (in green) and Bnab (in red), against DNA1. This graph is identical to Fig. 1B. **B.** Results from a replicate experiment using a different stock of beads, showing that the selectivities are reproduced except for the Bnab sequences (in red), which have a systematically higher selectivity. **C.** Similar to A, but when selecting for binding to the beads in absence of the DNA1 target. The top selectivities are from the Bnab sequences (in red), indicating that they bind to the beads, a finding consistent with the discrepancy between A and B. Here, the differences in selectivities are also coming from differences of selectivity during amplification (Fig. S11). Consistent with Fig. S11, the top Germ sequences (blue dots) have in absence of the DNA1 target the worst selectivities.

**Figure S13:**
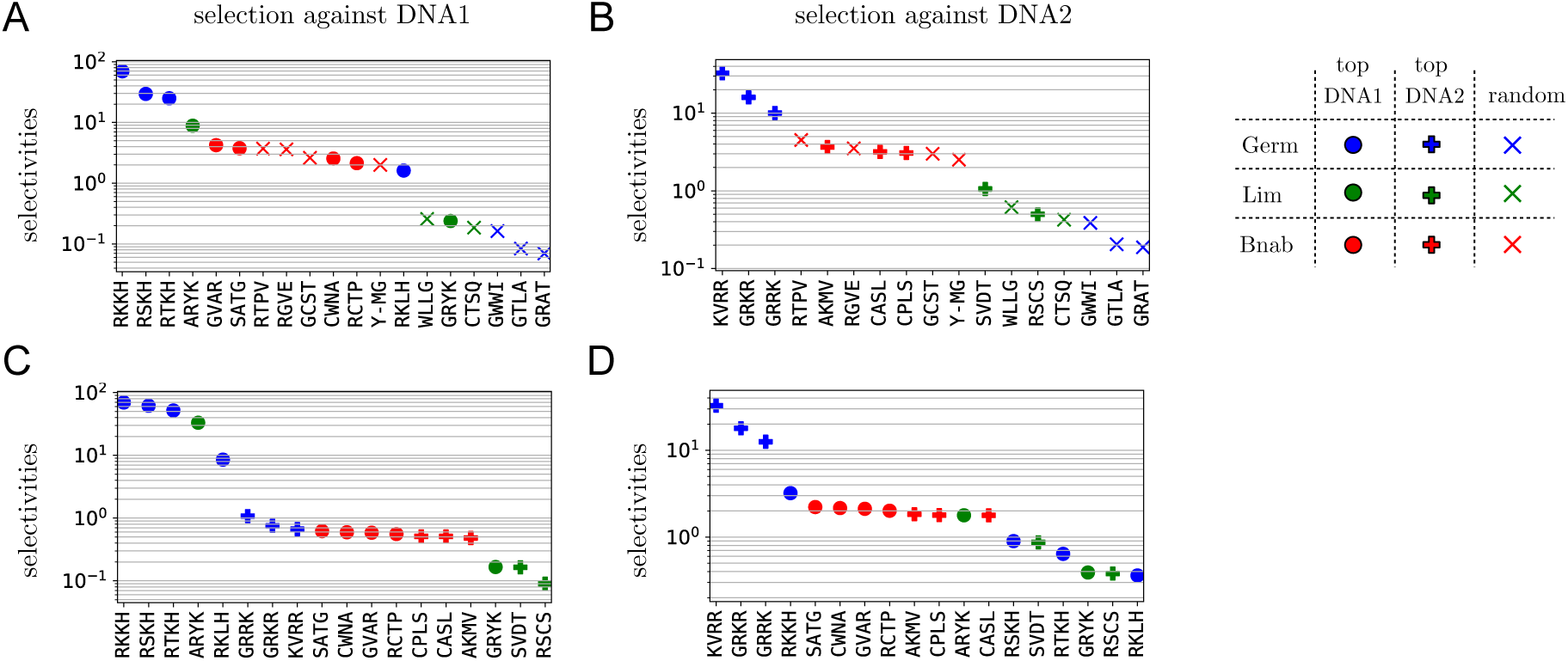
Cross selections with minimal libraries consisting of mixtures of top sequences against the DNA1 target (full circles) and top sequences against the DNA2 target (full crosses). **A**,**C.** Selection against the DNA1 target (same as Fig. 1B). **B**,**D.** Selection against the DNA2 target. The results confirm that some sequences from the Germ and Lim libraries bind specifically to the DNA1 target (blue dots and one of the green dots) and some sequences from the Germ library to the DNA2 target (blue crosses).

**Figure S14:**
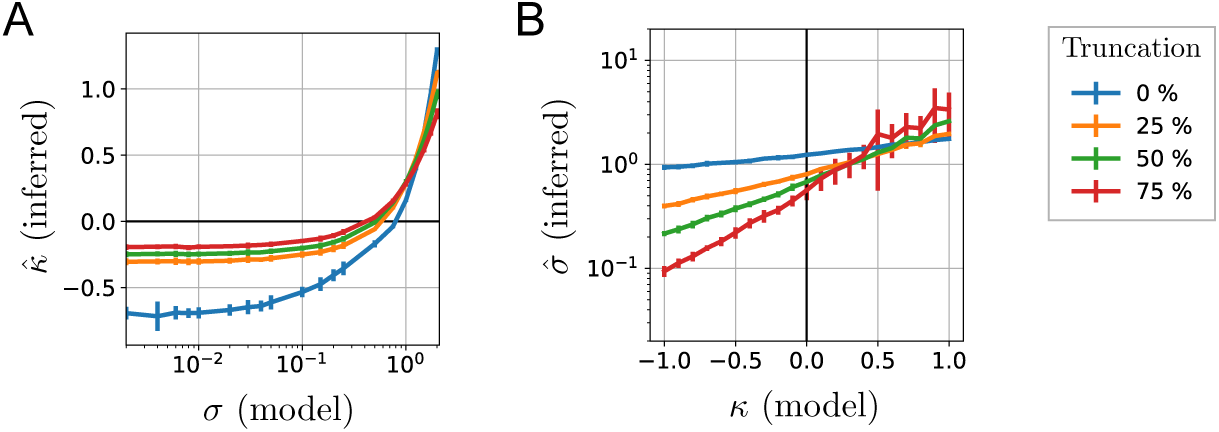
Relation between the parameter *σ* from log-normal fits and the parameter *κ*_*N*_ from generalized Pareto fits from numerical simulations. **A.** *N* = 10^4^ values were drawn from a log-normal distribution with parameters *µ* = 0 and varying *σ* (x-axis). The largest 25, 50, 75, 100 % of these values (i.e., 75, 50, 25, 0 % truncation) were fitted to a Pareto model with parameters *κ* and *τ*. The plot shows the estimation 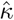 as a function of *σ*. Averages and standard deviations are taken over 25 independent realizations of the numerical experiment. It shows that limited sampling may cause a 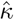 to be inferred from values drawn from a log-normal distribution when *σ* is small, here *σ <* 0.5. **B.** Inverse simulation: A truncated log-normal model is fitted to the largest 25, 50, 75, 100 % among 500 values (i.e., 75, 50, 25, 0 % truncation) drawn from a Pareto model with parameters *τ* = 0.115, *s*\* = 0.001 and varying *κ* (x-axis).

**Figure S15:**
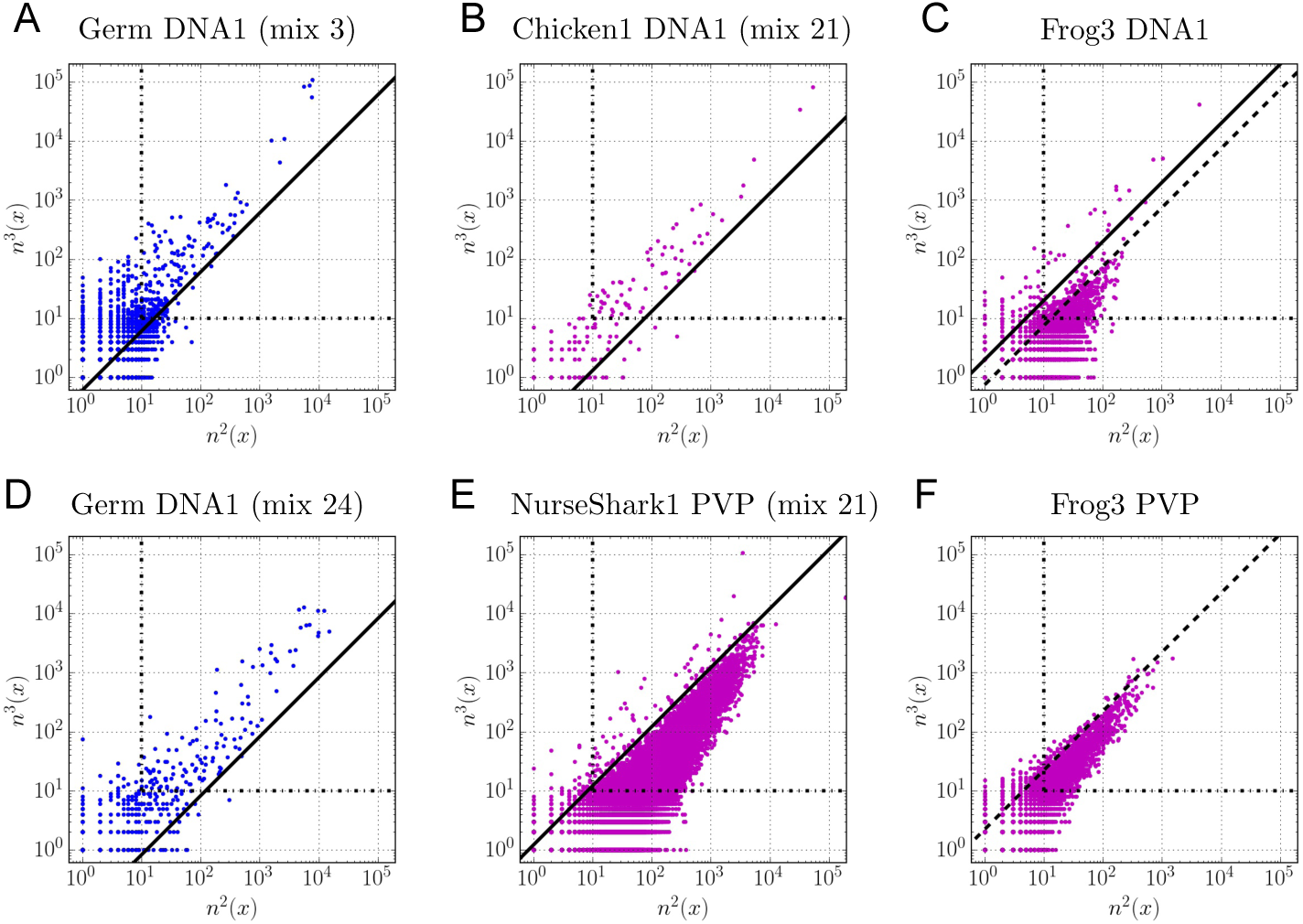
Definition of the threshold *s*\* above which selectivities *s* are considered for the experimental results reported here (A) and in Ref. [14] (B-F). As in Figure S3, the definition is based on a comparison between counts at the 2nd and 3rd cycles. The horizontal and vertical lines correspond to the criteria *n*^2^(*x*) ≥ 10 and *n*^3^(*x*) ≥ 10. The plain oblique line corresponds to the definition of *s*\* in this work. In the case of the selection of the Frog3 library against the DNA1 target, it differs from the value of *s*\* used in our previous work [14] (dotted oblique line) which failed to discard many selectivities coming from unspecific binding. In the case of the selection of the Frog3 library against the PVP target, all measured selectivities may be attributed to unspecific binding and we are therefore not including the inferred values of *σ* and *κ* in Fig. 4.

**Figure S16:**
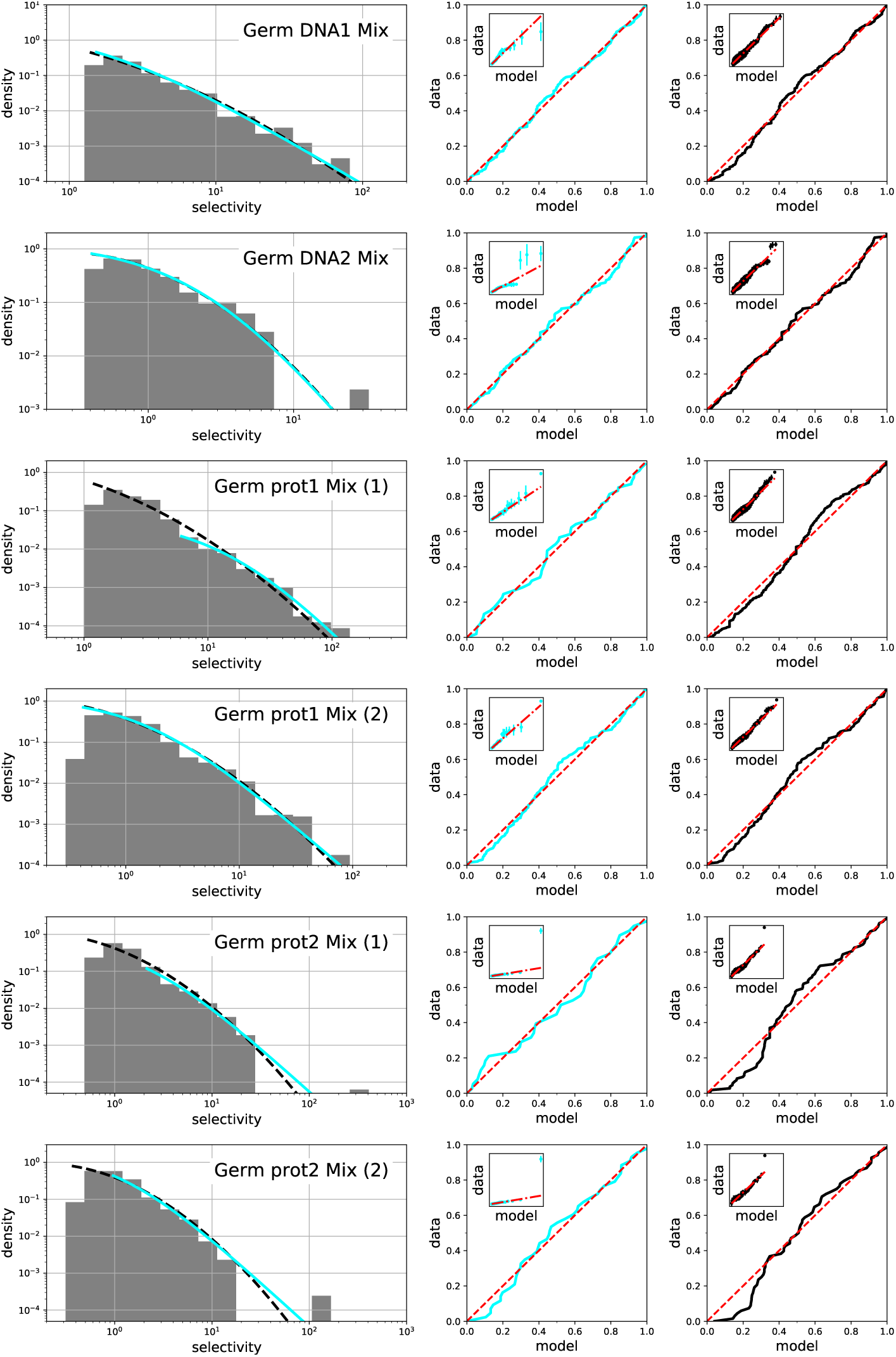
Assessments of the qualities of the fits of the selectivities to generalized Pareto distributions (cyan) and to log-normal distributions (black) for selections of the Germ library. The different graphs correspond to selections against different targets. For the protein targets prot1 and prot2, results from two replicate experiments are presented. All selectivities are computed by comparing the frequencies at the 2nd and 3rd cycle. The graphs on the right show the P-P and Q-Q (inset) plots for each fit. Perfect fits would correspond to the red dotted lines.

**Figure S17:**
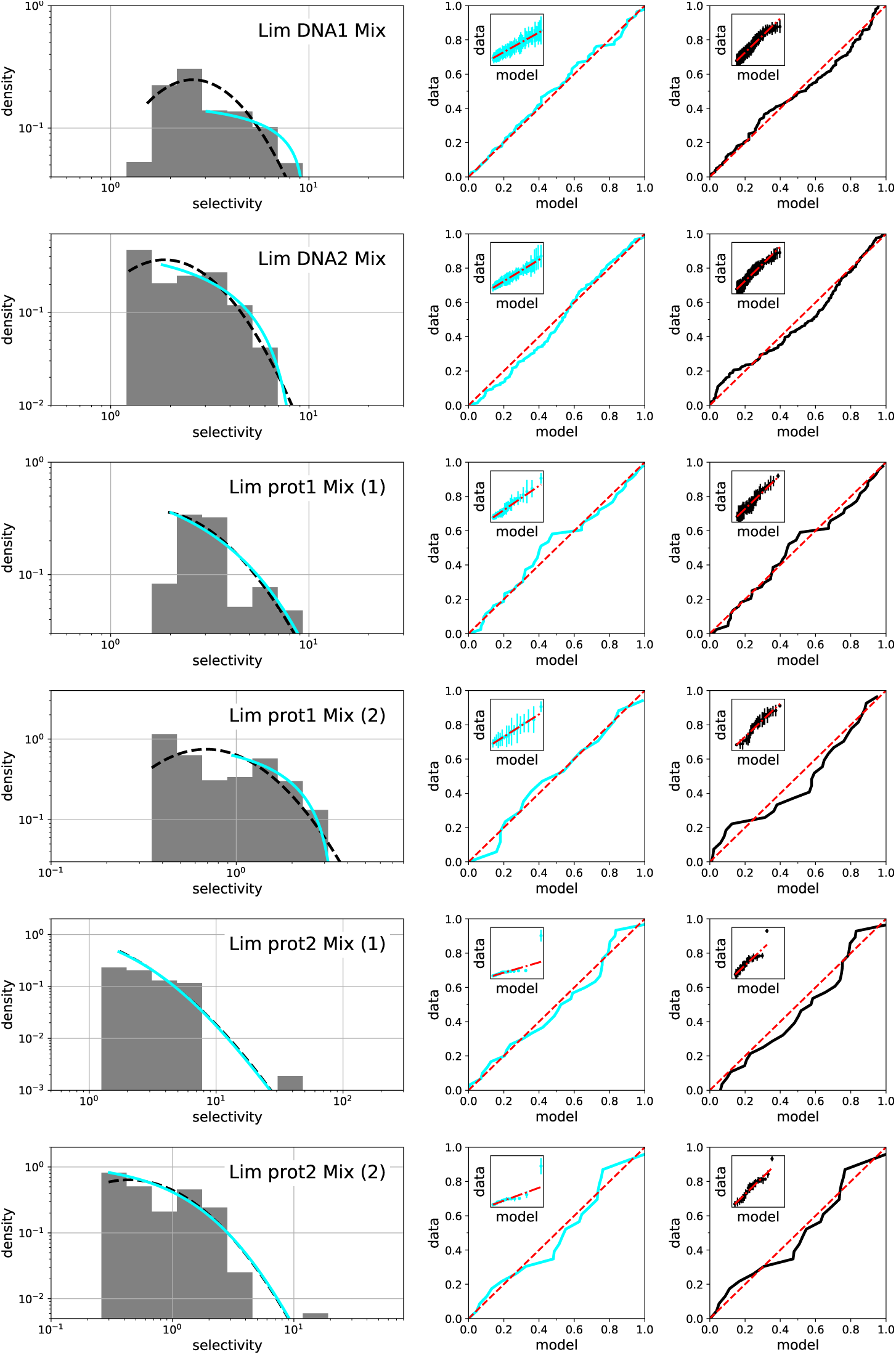
Same as Fig. S16 but for the Lim library instead of the Germ library.

**Figure S18:**
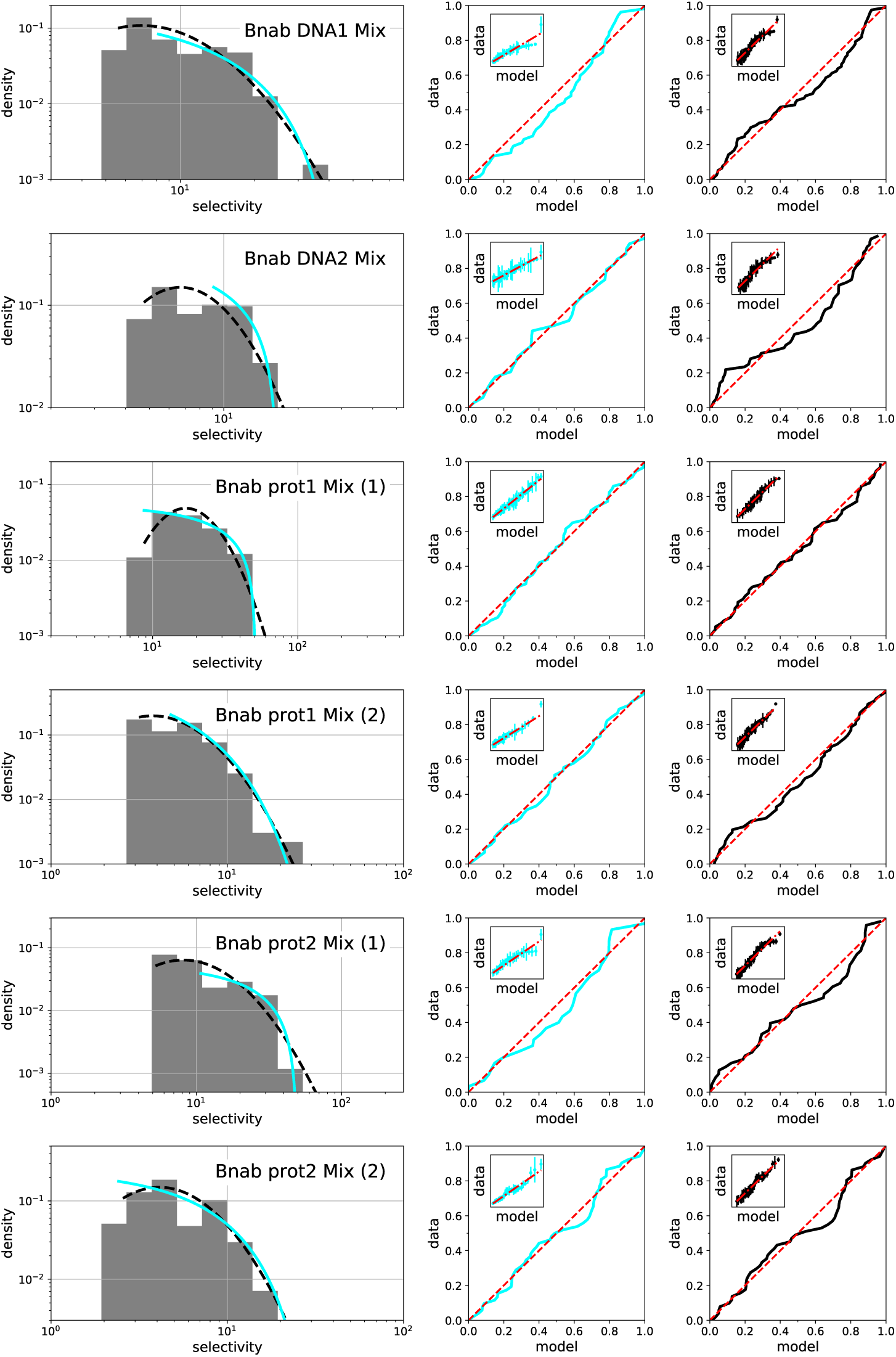
Same as Fig. S16 but for the Bnab library instead of the Germ library.

**Figure S19:**
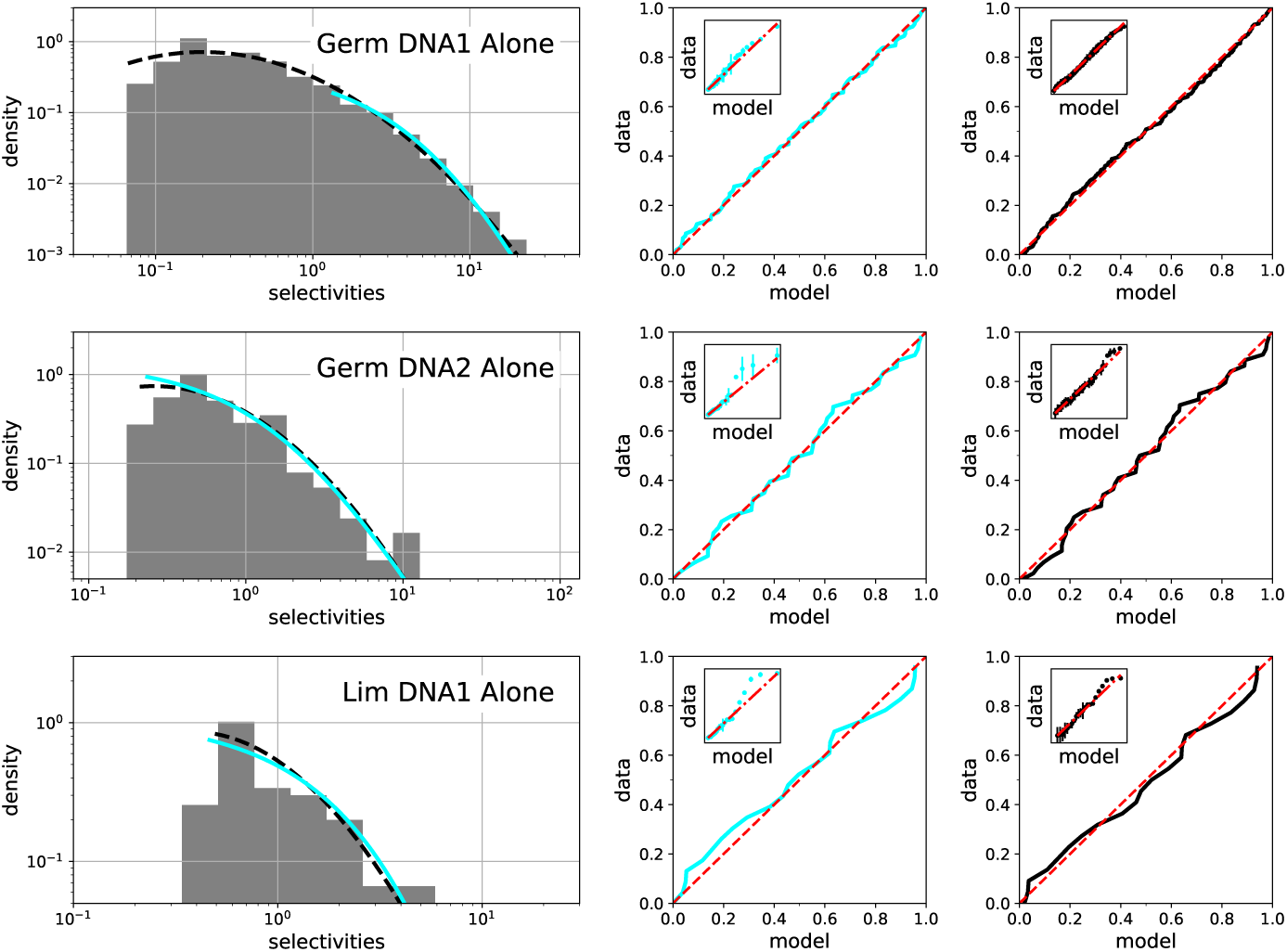
Same as Fig. S16 for the Germ library selected in isolation rather in a mixture with the two other libraries.

**Figure S20:**
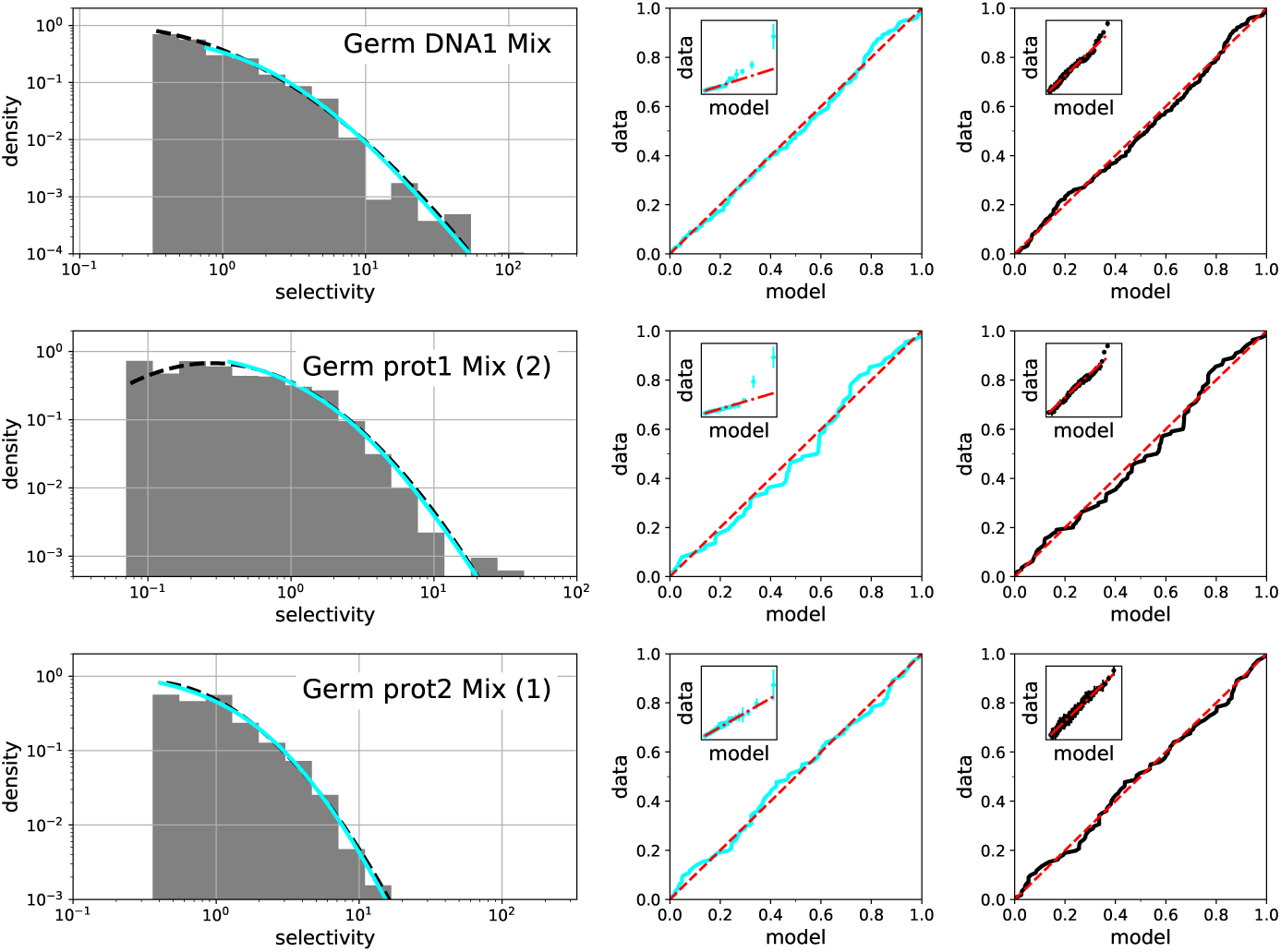
Same as Fig. S16 but for selectivities computed from a comparison between the 3rd and 4th cycle instead of the 2nd and 3rd cycle.

**Figure S21:**
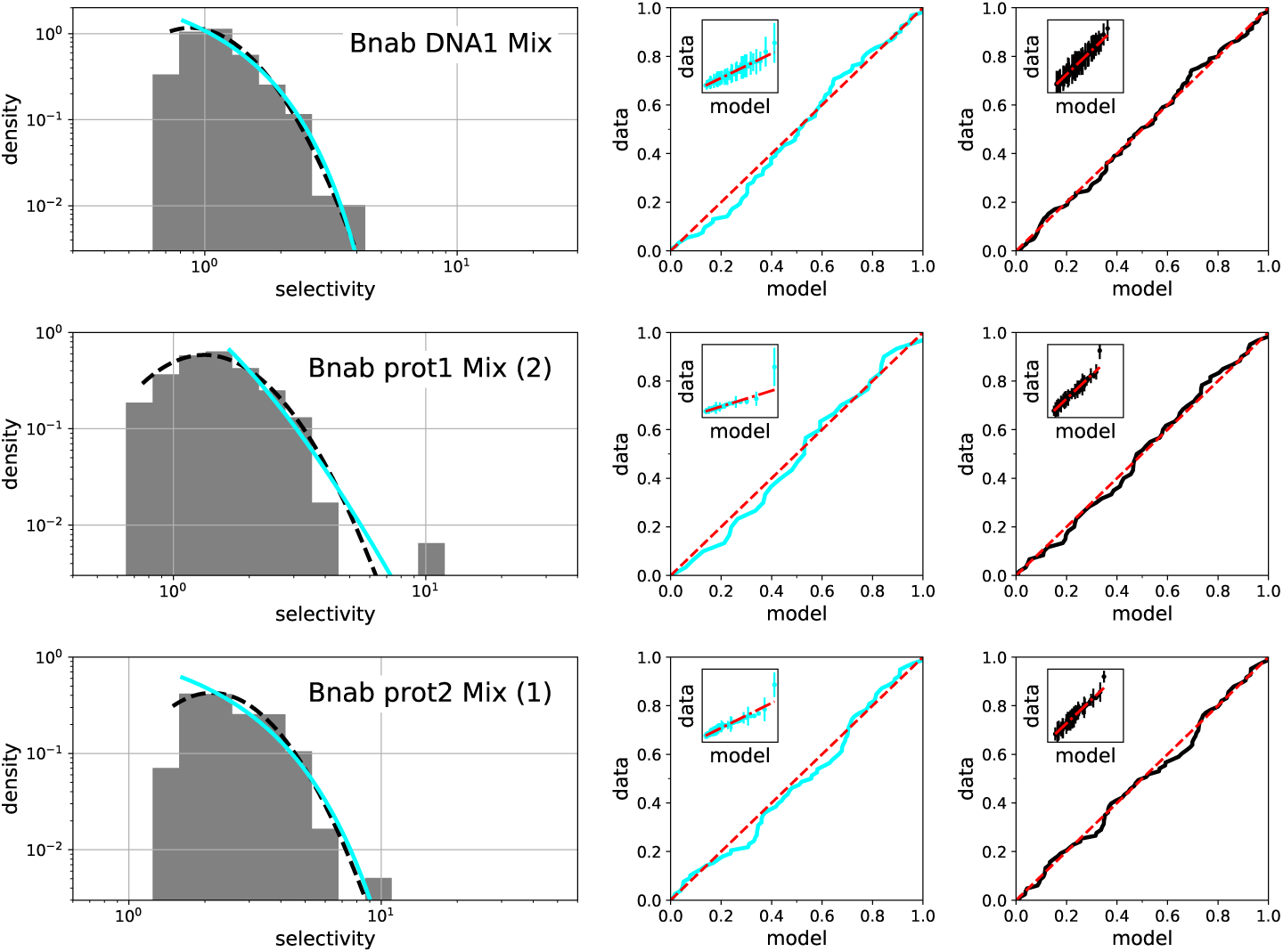
Same as Fig. S20 (selectivities computed from a comparison between the 3rd and 4th cycle) but for the Bnab library instead of the Germ library.

**Figure S22:**
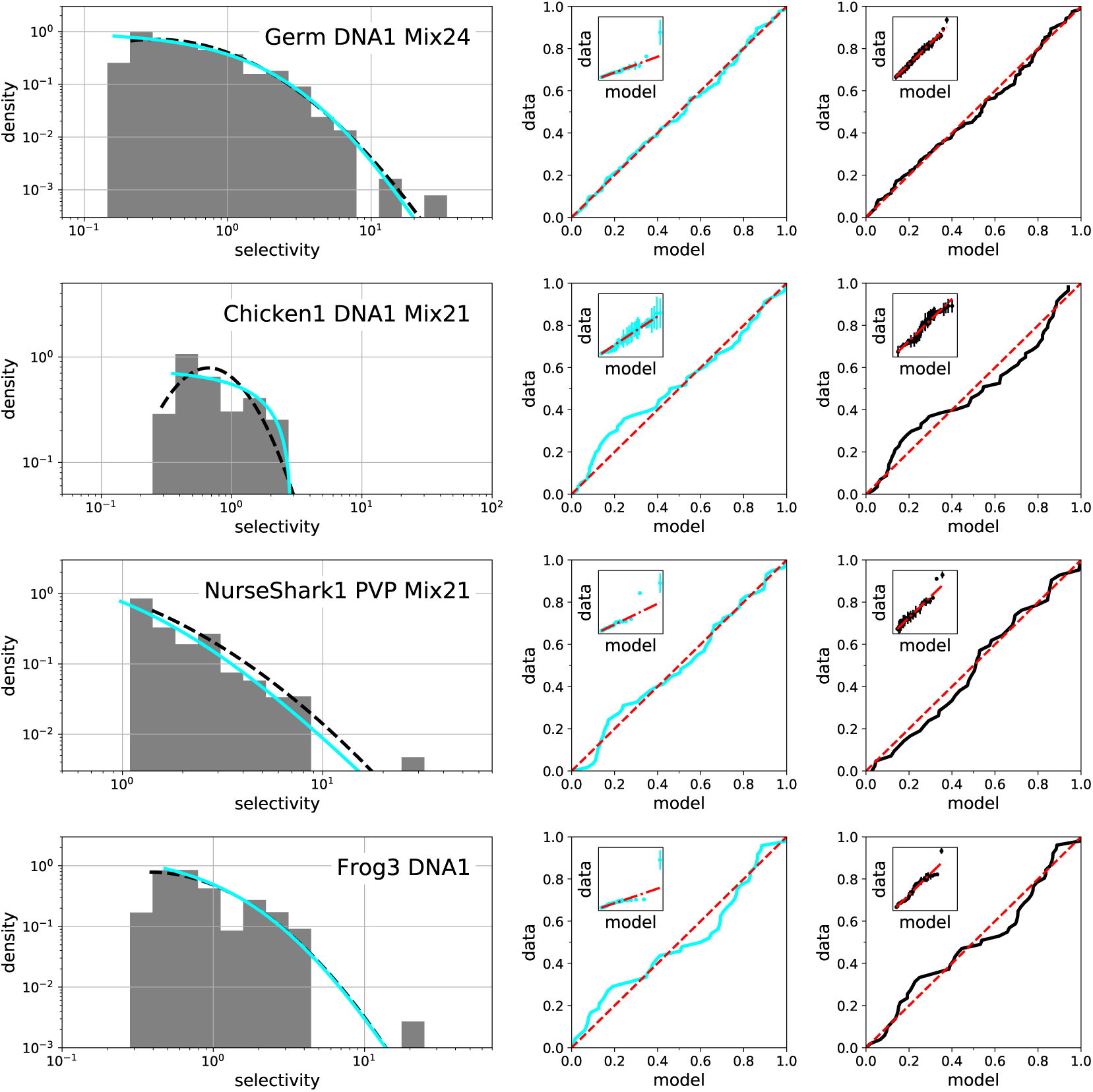
Same as Fig. S20 but for the experimental results reported in Ref. [14].

